# Probabilistic modelling improves relative dating from gene phylogenies

**DOI:** 10.1101/2024.09.30.615760

**Authors:** Moisès Bernabeu, Carmen Armero, Toni Gabaldón

**Affiliations:** Barcelona Supercomputing Centre (BSC-CNS). Plaça Eusebi Güell, 1-3, Barcelona, 08034, Spain; Institute for Research in Biomedicine (IRB Barcelona), The Barcelona Institute of Science and Technology, Baldiri Reixac, 10, Barcelona, 08028, Spain; Department of Statistics and Operations Research, Universitat de València, Burjassot, 46100, Spain; Catalan Institution for Research and Advanced Studies (ICREA), Barcelona, Spain; Centro de Investigación Biomédica En Red de Enfermedades Infecciosas (CIBERINFEC), Barcelona, Spain

**Keywords:** Phylogenomics, Relative dating, Bayesian Inference, Gene trees, Branch length distribution

## Abstract

1. Establishing the timing of past evolutionary events is a fundamental task in the reconstruction of the history of life. State-of-the-art molecular dating methods generally involve the reconstruction of a species tree from conserved, vertically evolving genes, and the assumption of a molecular clock calibrated with the fossil record. Although this approach is extremely useful, its use is limited to speciation events and does not account for genes following different evolutionary paths. Recently, an alternative methodology for the relative dating of evolutionary events has been proposed that considers the distribution of branch lengths across sets of gene trees.
2. Here, we validate this methodology by comparing the relative age estimates with a fossil-calibrated phylogeny and propose a model-based formalisation using a Bayesian framework.
3. Our analyses revealed that the normalisation of the distances of interest with the branch lengths of a reference clade present across the set of gene trees results in narrower distributions, allowing the correct inference of the relative ordering of evolutionary events.
4. We show that distributions of normalised lengths can be modelled using gamma or lognormal distributions and demonstrate that inference of the posterior distribution of the mode allows accurate relative age estimation, as assessed by a strong correlation with the molecular clock-dated tree.

## Introduction

Evolutionary biology aims at reconstructing the past history of living organisms. This process involves inferring a timeline, which minimally includes a relative ordering of events and, ideally, time estimates framed within the geological history of Earth (Reisz & Müller, 2004). The fossil record, coupled with radiometric dating and stratigraphy, can provide relatively accurate estimates of the time at which different groups of organisms lived, but its application is mostly limited to organisms containing fossilisable structures.

Molecular dating is a more recent dating approach that estimates divergence times between homologous sequences by leveraging the fact that they accumulate differences over time. This method relies on the concept of a molecular clock, which posits a correlation between evolutionary time and genetic sequence divergence (Zuckerkandl & Pauling, 1965). Importantly, the molecular clock can be calibrated using dated fossils, allowing the placement of molecular dating estimates within the geological timeline. Standard molecular dating typically begins by reconstructing species relationships from genetic data, focusing on orthologous genes (Boussau & Daubin, 2010; Kapli et al., 2020; Zuckerkandl & Pauling, 1965) and excluding genes showing signs of signal saturation or horizontal transfer (Philippe et al., 2011). The resulting phylogeny is then time-calibrated by assigning fossils to ancestral nodes, which constrains divergence times and allows the estimation of the molecular rate. This, in turn, enables the inference of divergence times for all speciation events in the tree (Dos Reis et al., 2016; Thorne et al., 1998).

Molecular dating has been successful in establishing the divergence of many macro-organismal lineages (Kumar et al., 2017). However, this method presents several important limitations. First, its accuracy heavily relies on the correct interpretation of the fossil record, which can lead to conflicting results across studies (Porter & Riedman, 2023). Second, a robust fossil record is lacking for most organisms in the Tree of Life, and many have no known fossils at all. Finally, molecular dating’s focus on reconstructed speciation events means it cannot precisely date evolutionary events that do not correspond to nodes within the phylogenetic tree.

To address some of these limitations and gain insights into a wider range of evolutionary events, an alternative framework using gene trees has been proposed: the branch length ratio method. This method infers relative timing by analyzing branch length ratios in gene trees (Pittis & Gabaldón, 2016a) and was initially applied to investigate gene acquisition in the lineage from the first eukaryotic common ancestor to the last common ancestor of extant eukaryotes (Pittis & Gabaldón, 2016a; Susko et al., 2021; Vosseberg et al., 2021).

This method is based on the premise that differences in evolutionary rates across gene trees can be taken into account by measuring ratios between a branch of interest and a “reference” tree distance that corresponds to the same divergence time in all gene trees. This reference tree distance is generally based on a well established evolutionary event such as the origin of a clade (the last common ancestor of eukaryotes in the studies above), that can be identified in all trees, and is considered as the median of the tip-to-event distances. As the clade has the same age in all gene trees, differences in the normalizing length would be informative on differences in rates. Hence, branch lengths normalized in this way in different gene trees are expected to represent relative differences in divergence time, and not in molecular rates. Doing so for an event of interest in a set of gene trees provides a distribution of relative ages with an associated uncertainty (variance).

Hence, the branch length ratio method provides a new framework for analysing the relative timing of evolutionary events which is independent of the fossil record and is not limited to speciation events. However, several caveats of this methodology have been discussed. First, the presence of unsampled or extinct (ghost) lineages may confound to some extent the branch length ratio analysis conclusions, particularly when applied to gene transfers (Susko et al., 2021; Tricou et al., 2022). However, simulations have shown that, even in the presence of ghost lineages, relative time inferences are overall more likely to be correct, and that the potentially conflicting lineages can be assessed (Bernabeu et al., 2024). Secondly, the modelling implemented by (Pittis & Gabaldón, 2016a) to identify distinct waves of gene acquisitions (a mixture of normal distributions) was criticised by showing that a lognormal distribution better fit the data (Martin et al., 2017). Although the conclusions of the paper were not dependent on this modelling (Pittis & Gabaldón, 2016b; Susko et al., 2021), the criticism underscored the lack of a thorough mathematical formalisation of the method.

Here, we developed a probabilistic framework for the branch length ratio method. To test our methodology, we inferred relative ages of clades from genome-wide sets of gene phylogenies and benchmarked them using a well-established molecular clock-dated tree of mammal evolution as the ground truth (Álvarez-Carretero et al., 2022). We found that both the gamma and lognormal distributions properly fit the empirical distributions of normalised branch lengths, which were generally skewed. Moreover, the use of a Bayesian framework allowed us to infer a posterior distribution for the modes as the best proxy for event timing, and perform a statistically-sound assessment of their relative ordering.

## Methods

### Sequence data

We selected a taxonomically-balanced set including 24 out of the 72 species considered in a recently reconstructed dated phylogeny of mammals (Álvarez-Carretero et al., 2022), and downloaded their corresponding genomes and gene annotations from Ensembl v101 (Cunningham et al., 2022) as of September 2022 (Table S1). Note that the selection of 24 out of the 72 species is necessary for the phylome approach (see below) as, to ensure accurate tree calculation in a reasonable time, a limit on the number of homologous sequences per tree is implemented.

### Phylome generation

We extracted the protein sequence of the longest isoform of each protein encoded in the selected genomes, and reconstructed a phylome (i.e., a complete collection of phylogenies from genes encoded in a genome of interest), using *Homo sapiens* as a seed, and using the PhylomeDB pipeline as implemented in phylomizer (https://github.com/Gabaldonlab/phylomizer) (Fuentes et al., 2022). In brief, for each *Homo sapiens* protein (seed), the pipeline runs a BLAST v2.13.0 (Altschul et al., 1990) against all 24 selected species’ proteomes. We selected those hits with an alignment coverage over the query sequence higher than 33% and an e-value lower than 1e-5. In addition, we limited the homologous gene set to the top 200 sequences. The sequences for each gene family were aligned using MUSCLE v3.8.1551 (Edgar, 2004), MAFFT v7.407 (Katoh & Standley, 2013) and Kalign v2.04 (Lassmann & Sonnhammer, 2005) in forward and reverse orientation. Then, the six resulting alignments were merged into a consensus alignment using M-Coffee v12.0 (Wallace et al., 2006). The consensus alignment was trimmed using trimAl v1.4.15 (Capella-Gutierrez et al., 2009) with a gap threshold of 0.1 and conserving a minimum of 30% of the positions of the original alignment. This trimmed alignment was used to reconstruct a phylogeny using IQ-TREE v1.6.9 (Nguyen et al., 2015), under the best-fitting model selected from a subset of the available ones (DCmut, JTTDCMut, LG, WAG, VT) using ModelFinder (Kalyaanamoorthy et al., 2017), and, the support was assessed using 1,000 ultra-fast bootstrap replicates. Using 1000 randomly-chosen trees we tested the impact of using more complex models (LG+C60+G4, LG4X or LG4M) and concluded this was minimal for our dataset (see Supplementary Results, Fig. S18).

### Tree distance calculation

We implemented a custom script for calculating the phylogenetic distances of interest (Fig. 1). This script takes as input a gene tree, the rooted reference species tree (in this case, the pruned dated tree from (Álvarez-Carretero et al., 2022)), and a table listing the ancestral events of interest (the origin of primates, boreoeutherians, therians and placental mammals) with the species descending from it. From the species tree, the script derives two types of information (Fig. 1a, 1): (i) a “species-to-age” dictionary, numbering all nodes ancestral to the seed sequence (from 1, the most recent to n, the most ancient) and indicating, for each other species in the tree the most recent common ancestor (MRCA) with respect to the seed species and (ii) a “first split” dictionary, defining for each considered event the two descendant clades (Fig. 1a, 1). Note that the input species tree does not need to be dated, as only topological information is used.

**Fig. 1.**
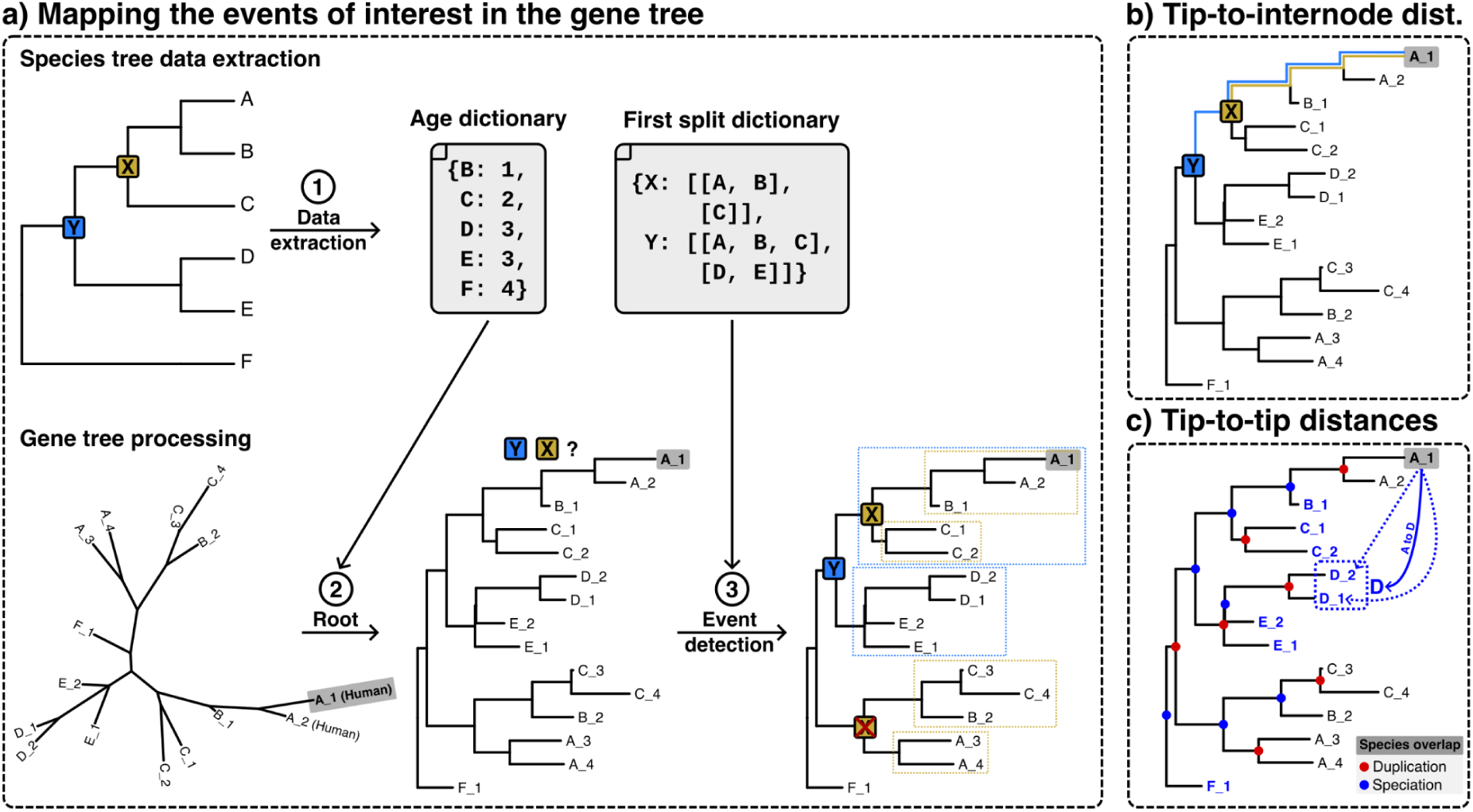
Schematic representation of the computational pipeline used for obtaining evolutionary distances. A1 (in bold and shadowed) represents the seed sequence. a) (1) Extraction of the age and first split dictionaries from the species tree, (2) gene tree rooting using the age dictionary, and (3) detection of the events’ MRCA by assessing the presence of the first split, boxes in the tree mean the two monophyletic groups after the first split for each event in the tree, the X event below cannot be considered, for two reasons i) it does not contain the seed, and ii) despite all its descendants belonging to the X group, its first split does not match the species tree one. b) Tip-to-internode distance calculation, once the event’s node is detected, the distance between the seed and the node is calculated by summing up the branch lengths in their connecting path. c) The tip-to-tip distance is calculated by obtaining the distance between the seed tip and all its orthologs (in blue) in each of the other species. In the example, the distance of the seed (A1) to the species D is calculated by retrieving the distances to all the co-orthologs belonging to the species D (D1, and D2, indicated with the blue dashed box).

Each gene tree is rooted at its oldest node using the species-to-age dictionary, as described in (Huerta-Cepas et al., 2007), (Fig. 1a, 2). Then, duplication and speciation nodes are inferred using the species overlap algorithm (Huerta-Cepas et al., 2007), as implemented in the ETE3 Python package (Huerta-Cepas et al., 2016). In addition, the pipeline uses a taxonomy table to label the tree leaves to indicate to which clade they belong (Fig. 1a, 3). We retrieved the subtrees of the events of interest, that is, those monophyletic clades whose MRCA is the event of interest. To this end, we designed an “MRCA function”, which retrieves the largest monophyletic subtree containing all the sequences from the species belonging to the target clade that accomplish the following conditions: (i) any MRCA to tip distance is 0 (this occurred in a few cases where the sequences descending from the MRCA are identical due to alignment trimming), (ii) the first split in the subtree is congruent with the species tree (for instance, to detect the boreoeutherian ancestor, we require the first split to divide primates and rodents and lagomorphs), allowing for missing species; and (iii) the subtree contains the seed sequence (Fig. 1a, 3). Note that gene trees where the relevant MRCA are not present, due to gene-tree vs species-tree incongruences will be discarded and not considered. We calculated two types of distances, first, tip-to-tip distances, which are the distances from the seed sequence to all the other tips in the tree that are orthologous to the seed (e.g., from humans to mice). Orthology was inferred using the species overlap algorithm implemented in the ETE3 package (Huerta-Cepas et al., 2007). Second, the tip-to-internode distances, which are the distance between the seed and a speciation node corresponding to an internal node (one of the events considered or ancestor of the human lineage).

### Normalisation of tree distances

To account for across-gene differences in evolutionary rates, we used a phylogenetic normalisation approach similar to that of (Pittis & Gabaldón, 2016a). Here, we used Primates as the reference clade for normalisation. We have chosen this group as it is close enough to the seed species (*H. sapiens*), and large enough to allow sampling a significant number of branch lengths. We thus normalised the raw distances of interest by dividing them by the median of the MRCA-to-tip distances of the Primates clade. We observed some large distances due to (principally) small normalising groups (in terms of number of tips or short branch lengths), which provided near to 0 normalising factors and then extremely large normalised distances. To solve this, we removed normalised distances greater than the 99th quantile for the tip-to-internode distances and the 90th quantile for the tip-to-tip distances (we used a more stringent quantile in tip-to-tip distances as we observed extremely long normalised distances caused by the effect of some gene family expansions). All the tree functions and calculations were implemented using ETE3 (Huerta-Cepas et al., 2016).

### Modelling tree distance distributions

Normalised evolutionary distances are positive real numbers that exhibit a right-skewed distribution. We chose the gamma and the lognormal distribution as data-generating models, as both of them are highly tuned to the shape exhibited by the data and have analytical expressions for their most important features, especially the mode, which will be the characteristic used for the relative dating. The continuous gamma distribution *Ga*(α, β) depends on two parameters, shape α > 0 and rate β > 0. Its mean and variance are α/β and α/β², respectively. The mode is 0 if α < 1 and (α − 1)/β otherwise. The continuous lognormal distribution *logN*(µ, σ²) depends on two parameters, − ∞ < µ < ∞ and σ > 0. Its mean and variance are exp(µ + σ²/2) and ((exp{σ²} − 1) exp{2µ + σ²}), respectively. The mode is exp{µ − σ²}.

Our methodological statistical framework is Bayesian Inference (BI), which we will use to infer the parameters of the two presented distributions. The three essential elements of a Bayesian statistical analysis are: first, a prior probabilistic distribution π(θ) for all the parameters of interest θ. In our case, θ = (α, β) when dealing with the gamma distribution, and θ = (µ, σ) in the case of the lognormal model. Second, the likelihood function *L*(θ) of the parameters for the observed data, which we will represent as *D* from now on. The likelihood is the product of the densities across the normalised distances using the parameters set θ, resulting in the probability of observing our data given θ. In our study, the data are the normalised distances. And finally, the posterior distribution for θ, π(θ|*D*), which combines the prior and the data information using Bayes’ theorem as follows

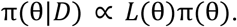

We decided to give more importance to the data than to prior distributions, and therefore considered prior independence and selected wide and poorly informative uniform distributions *U*(0, 100) for (α, β, σ), and *U*(− 100, 100) for µ.

The subsequent posterior distribution of the parameters of both the gamma and the lognormal distributions is not analytical. For this reason, we use Markov Chain Monte Carlo (MCMC) methods, in particular Gibbs sampling, to obtain an approximate sample of this distribution to allow us to make inference about the target parameters and the derived output quantities. MCMC was implemented via JAGS v4.3.0 (Plummer 2003), using default parameters. We ran three independent chains with 100,000 iterations each, removed 10% of the initial iterations which we considered a burn-in period, and used a thinning of ten iterations. Convergence was assessed using both graphic and numerical diagnostic tools. In particular, the Gelman-Rubin’s statistic, which is the quotient of the variances of the chains within and between the independent runs, with values close to 1 indicating convergence, and the effective sample size (ESS) which accounts for the number of independent samples. The greater the ESS, the more samples are suitable to behave as the posterior. In addition, we plotted the traces and autocorrelation for all the parameters and chains. The runtime of the Bayesian inference in our largest set (primates, with 4,335 trees) was 1h 30 min 56 s in an x86-64 machine based on Intel® Xeon® Platinum 8160.

To assess the robustness and sensitivity of this procedure, we collected random subsamples from the original tree sets for each considered event. For instance, the boreoeutherians’ event node is present in a set of trees, *B*. We randomly subsampled trees (*b_*i*_* ⊂ *B*) and repeated the inference process for both models in these subsets. The size of the subsamples was set to represent incremental percentages of the total sample size, obtaining subsets from 10% (*b*_10%_) to 100% (*B*) including all 10% increments, plus 15% and 25% to have higher resolution in the 10-30% range.

## Results

### Genome-wide distribution of phylogenetic distances

We first set out to investigate the shape of the distribution of phylogenetic distances obtained from a genome-wide collection of gene phylogenies (i.e., a phylome, Sicheritz-Pontén & Andersson, 2001). For this, we reconstructed the human phylome in the context of 24 mammalian species, for which a recent highly resolved timed phylogeny is available ((Álvarez-Carretero et al., 2022), see Methods). This phylome includes 16,828 gene trees and is available for browsing or download at PhylomeDB with the PhylomeID 0593 (Fuentes et al., 2022). We first measured, for each gene tree, the tip-to-internode phylogenetic distance between the human seed gene and four events of interest: namely the origin of primates, boreoeutherians, placentals, and therians.

All resulting tip-to-internode distances distributions were close to 0 and largely overlapped (Fig. 2a). As previously done by Pittis and Gabaldón (2016a), and to account for differences in evolutionary rates across gene families, we normalised the raw distances by dividing them by the median of the branch lengths observed in the primates clade (see Methods). This normalisation resulted in sharper distributions that are farther away from 0 and are more separated among them (Fig. 2b). This allows a better relative timing of the considered events based on the ordering of the peaks of these normalised distance distributions. This ordering agrees with the sorting of the same events according to the dated species tree. As expected from an erosion of the phylogenetic signal with time, distances to older events were associated with a higher dispersion, and with a lower number of gene trees containing that event (Fig. S1d). Note that estimated relative age peaking sharply around 1 for primates underscores the suitability of the use of the median of branch lengths of the primate branches as a normalising factor.

**Fig. 2.**
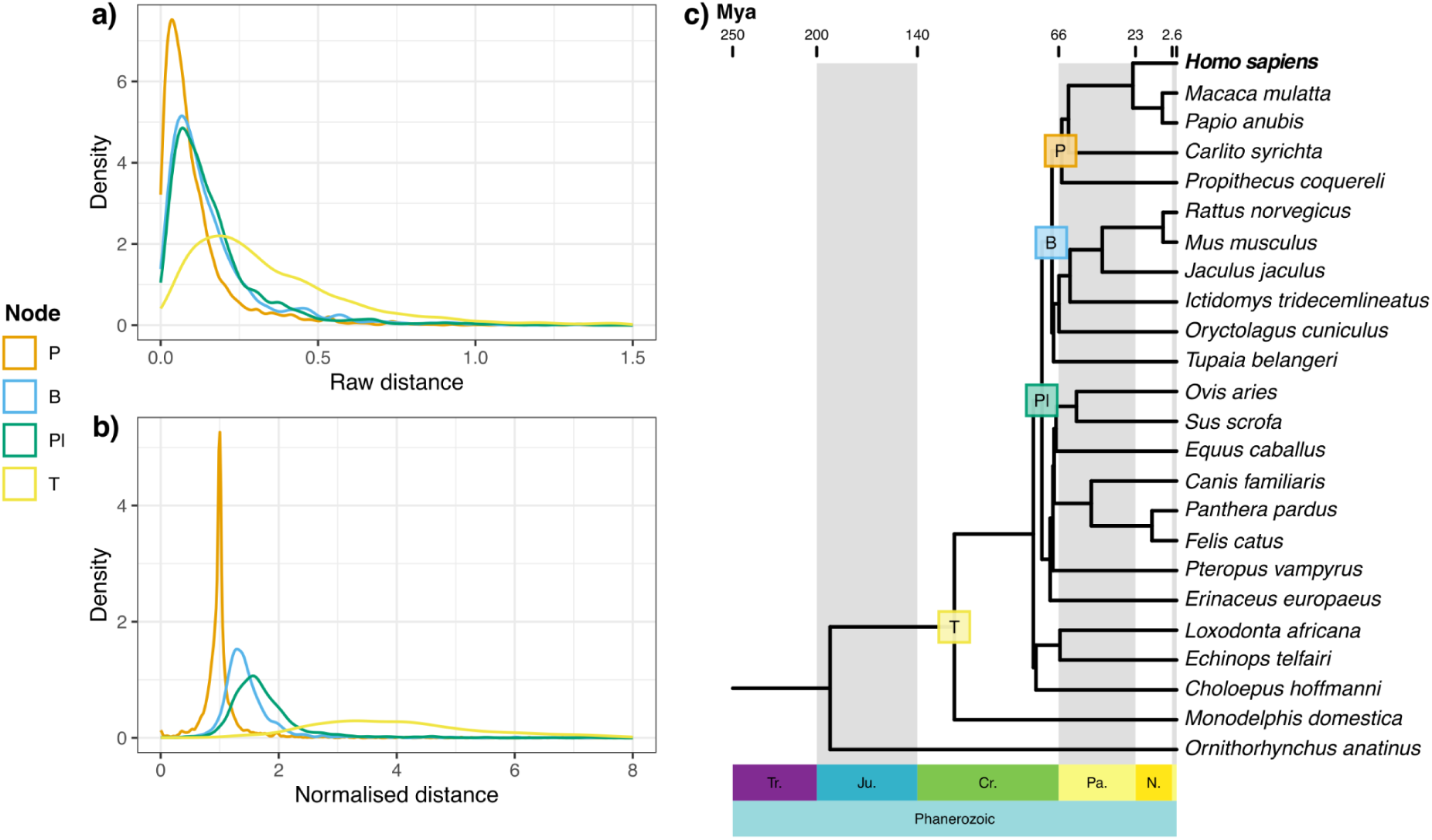
Distributions of the tip-to-internode distances to the events of interest. a) Distribution of tip-to-internode raw distances. b) Normalised tip-to-internode distributions for each event. c) Dated species tree from (Álvarez-Carretero et al., 2022), labelled nodes are those assessed in the distribution of a) and b). The legend shows the group that originated in the event, T: Theria, Pl: Placentalia, B: Boreoeutheria and P: Primates. Empirical cumulative distribution functions for the distributions in a and b are available in Fig. S16.

To assess the impact of using a different normalising clade, unrelated to the seed of the tree, we computed tip-to-internode distances normalising with the Laurasiatheria and the Rodentia clade. The distances obtained with either of these groups or primates as normalising clades were significantly correlated, and resulted in identical conclusions in downstream analysis regarding the relative ordering of events (Supplementary Results, and Fig. S19).

We used the same approach to calculate tip-to-tip distances between the seed human sequence of each tree and its orthologs in each of the other species (see Methods). In this case, raw distances (Fig. S2) show values in a restricted range around 0 and 2, while normalised distances (Fig. S3) had a wider range, from 0 to 15, and were more separated and easier to discriminate. Unexpectedly, we found that the distances to the closest species to the human seed (*Papio anubis* –PAPAN– and *Macaca mulatta* –MACMU) had bimodal distributions. Upon further investigation, we found that gene trees underlying the first and second peaks differed in the encoded functions: the first peak (including genes having shorter normalised distances) contained mostly informational genes (DNA and RNA processing), whereas genes underlying the second peak (having longer normalised distances) are enriched in metabolic functions (Supplementary Methods 1.2 and Figs S4-S7). Another potential factor affecting these branch lengths is incomplete lineage sorting, which is pervasive in primates (Rivas-González et al., 2023). We kept both peaks and focused on the mode, the first peak in this case, considering the second peak as a source of variability rather than a source of information for the timing.

From these analyses, we conclude that the normalisation of tip-to-internode and tip-to-tip phylogenetic distances from collections of gene phylogenies has the potential to infer a correct relative timing of evolutionary events, as proposed earlier (Pittis & Gabaldón, 2016b; Susko et al., 2021). We also note that rate differences and incomplete lineage sorting may result in multimodal distributions after normalisation, particularly in recent events. For comparison, we explored alternative normalisation approaches, but concluded that they did not provide significant advantages over this one (see Supplementary Results).

We further explored the effect of normalisation using simulated trees. We observe that normalisation produces the same effects in simulated data, narrowing the distributions and separating the modes of each event. However, as compared to real gene trees, simulated data provided lower resolution to separate the different events from (Supplementary Results, and Fig. S20).

### Modelling the distribution of genome-wide phylogenetic distances

To infer the underlying probabilistic distribution of the branch lengths, we used gamma and lognormal models. We carried out Bayesian inference (BI) on the parameters of these distributions using MCMC sampling via JAGS (Plummer, 2003). We set non-informative prior distributions as the prior for each parameter of the gamma distribution, π(α) = π(β) = *U*(0, 100), as well as for each parameter of the lognormal distribution, π(µ) = *U*(− 100, 100), and π(σ) = *U*(0, 100). We ran three independent MCMC chains with 100,000 iterations each, removed 10% of the initial iterations which we considered as a burn-in period, and used a thinning of ten iterations. We further checked the convergence and autocorrelation using both numerical and graphical methods. We repeated the inference process in randomly generated subsamples of the total tree population for each studied event.

For the models and parameters tested, we observed convergence of the three independent chains, which provided enough posterior samples to infer the posterior distribution of the parameters. In the case of the gamma distribution, *R̂* values are close to 1 meaning that the within and between chains variability is similar. Moreover, the ESS values range from ∼6,152 to ∼95,527, these high ESS values allow us to treat the posterior samples as independent. Despite all the parameters converging, the autocorrelation decreases slowly (mean autocorrelation at lag 3 of 0.35); however, it reaches 0 in some lags. Regarding the lognormal model, all the *R̂* values are close to 1, as in the gamma model. The ESS ranges from ∼85,596 to ∼96,485, providing more independent posterior samples than the gamma model. The lognormal model improves the autocorrelation, which reaches a mean value of 0.003 in just 3 lags (Tables S2-S4). Both models accurately fitted the branch lengths. Moreover, the highest posterior density intervals are narrow (Table S5).

### The mode of the inferred Gamma distribution of branch lengths as a proxy for relative timing

We next explored the potential of model-based inference for the relative timing of evolutionary events using sets of gene trees. Evolutionary events (such as the origin of a new clade) are usually assumed to correspond to a single discrete point in time. Then, the observed variability in normalised branch lengths should result from analytical (alignment or phylogenetic inference errors) or biological factors (varying evolutionary rates not captured by the normalisation). Given the non-symmetrical nature of the distance distributions, we hypothesised that the mode of the distribution is the best proxy for the time point of interest, and tested this assumption by comparing the inferred modes with the corresponding distances in the dated tree (Álvarez-Carretero et al., 2022). The posterior distributions of the modes for each event (Fig. 4a) had very low dispersion, which means that different events can be easily distinguished.

To test whether the inferred modes were accurately retrieving temporal information from gene trees, we compared them with the molecular clock-dated species tree of our set of species (Álvarez-Carretero et al., 2022). We obtained the distances in Million years (My) from the *H. sapiens* tip to all the internal nodes in the path from the tip to the root in the dated species tree, and the same normalised distances for the phylome set of gene trees. Then we calculated the posterior distribution of the mode for each node. We also used the subsequent posterior distribution of the tip-to-tip distance and the corresponding distances in the species tree. The correlation between both dating methodologies was high (Fig. 3). Regarding the tip-to-tip distances (Fig. 3a), there are several species with equal distances, this is expected for the ultrametric property of the dated species tree, which means that the distance to all members of a monophyletic sister clade will be the same. Despite this, the inferred normalised distances agree with those in the dated tree. This correlation is even higher when focusing on the distances within the lineage leading to humans (Fig. 3b). These results indicate that the normalised distances obtained from collections of gene trees are a good proxy of time, and that they allow a correct sorting of evolutionary events.

**Fig. 3.**
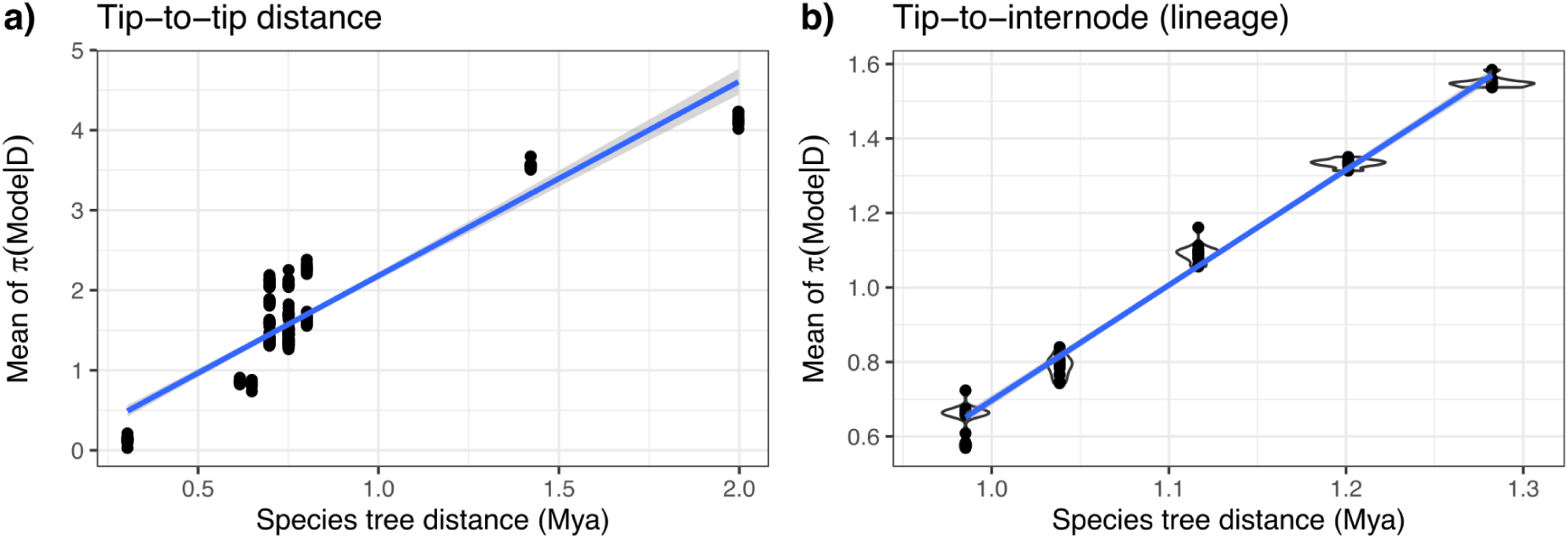
Correlation between the mean of the posterior sample for the mode and the species tree distance. a) Tip-to-tip distance *R*² = 0. 91 and b) tip-to-internode (human lineage except the human speciation and the root) distance *R*² = 0. 99.

### A probabilistic framework for relative timing

In Bayesian inference, we can compare two parameters, for example θ_1_ and θ_2_, through the subtraction or division of their posterior distributions. In the case of the subtraction, which we used here, the posterior distribution would be π(θ_1_− θ_2_|*D*), where θ*_i_* is the mode of the evolutionary event *i*. Thus, posterior distributions for the subtraction of two events (π(θ_1_ – θ_2_ |*D*)) around zero, would indicate no relevant difference between θ_1_ and θ_2_, and the most likely hypothesis is that they happened simultaneously. Conversely, posterior distributions for the subtraction far from 0 would refer to events that happened at different times.

We assessed whether this method accurately distinguished contiguous evolutionary events (Fig. 4b). All the comparisons between the timing of the events concluded that they occurred at different times with a probability of 1, although some of the compared clades occurred close in evolutionary absolute time (e.g., the origin of Placentalia and Boreoeutheria).

**Fig. 4.**
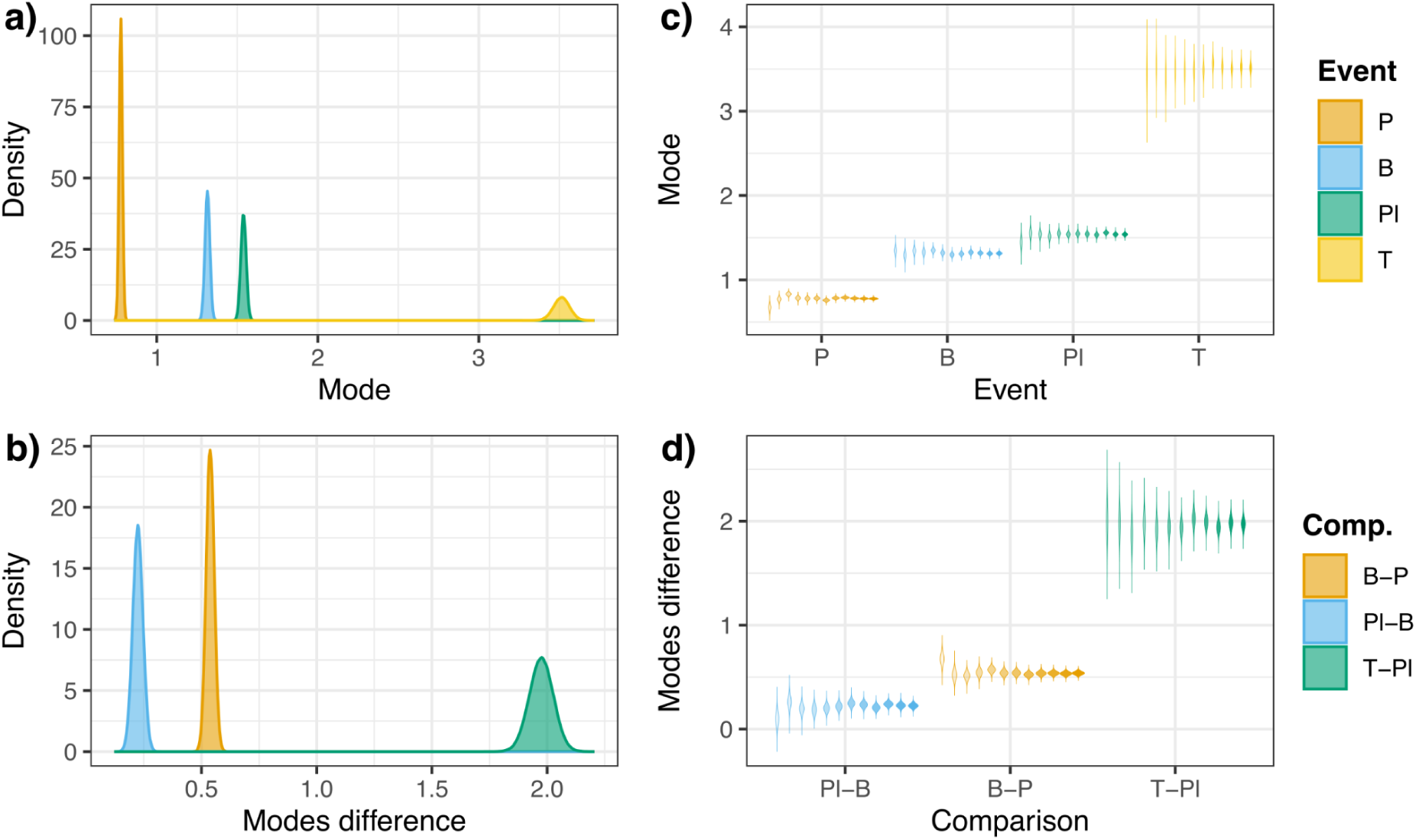
Posterior distributions of the events relative timing. a) Posterior distribution of the mode for each event. b) Posterior distributions of the difference between modes of adjacent events (Comp.: comparison groups). c) Posterior distributions, shown as violin plots, of the mode of each event for each subsampling set. d) Posterior distributions, shown as violin plots, of the difference between modes of adjacent events for each subsampling set. Subsampling ranges from 10% of the total number of trees to 100%, increasing by 10% each, we also included 15% and 25% of the trees. The more opacity of the violin, the higher the percentage of trees used. T: Theria, Pl: Placentalia, B: Boreoeutheria and P: Primates. Empirical cumulative distribution functions for the distributions in c and d are available in Fig. S17.

To assess the robustness of the approach, we repeated the inference on randomly chosen subsamples of the trees (Fig. 4c). The modes inferred from the subsamples or the full-dataset were always similar, with deviations ranging from 0.69% in the Boreoeutheria node to 10.47% in the primates node for the smaller subset. In this study, we used a seed-based approach, in which homologs are inferred by searching the seed sequence in a given set of genomes. Because of this homology search strategy, the events closer to the seed sequence are over-represented with respect to the ancestral ones. As a result of this, the subsampling has a stronger effect on the variability of the deeper nodes, as seen in Fig. 4c. Despite the congruence with the inferred mode, the standard deviation of the posterior distributions for the mode ranges between 0.005 in the primates node (most recent event) using the complete tree set to 0.081 in therians (deepest event) using 10% of trees (Figs S9 and S10a). As expected, the standard deviation of the inference increased when we used fewer trees, as there is less information to infer the statistic (Table S6). Importantly, however, the standard deviation values are small relative to the distance between modes and thus, such variation is unlikely to cause a shift in the relative timing (Fig. 4d). Furthermore, the standard deviation does not decrease constantly, there are some points in which the standard deviation of the posterior increases for a higher number of trees (25% for Theria and 20% for primates subsample in Fig. S10a).

This suggests that the nature of the gene trees in the subsample, and not just the number of trees, is important. We used the same subsampling strategy to assess whether, when using fewer trees, the method allows us to discriminate between events (Fig. 4d). The subtraction of posterior samples using less data showed that, despite using only 10% of the trees, the studied events could be differentiated with probability 1. As we observed in the events’ case, the variation between the standard deviation of the subsamples is around 70% (Table S7). However, the farthest comparison (Theria origin against Placentalia) shows a slightly higher standard deviation (Fig. S11a). The deviation from inferred differences between modes (i.e., using all the trees) is between 1% and 56% (Table S7, the highest values occur as a consequence of the small difference values, a small variation turns into a higher percentage, for instance, the difference between the 10% set and the full set of trees for the Placentalia and Boeroeutheria –56.41% deviation– is just 0.12). Despite this, the measure of interest here is the area under the curve of the modes’ comparison (*P*(*mode*_1_ – *mode*_2_> 0 | *D*)) when it is greater than 0, for the closest event we found that using the 10% of the trees the probability that Placentalia clade originated before Boreoeutheria clade is ∼1. Despite variations in the dispersion of the distributions, the differences between the inferred mode from different subsamples are low for both the events and the comparisons. This test supports the robustness of the inference method to provide probabilistic information about the sorting of evolutionary events.

Finally, to assess the robustness of the method when comparing two clades that do not include the human lineage, we compared the relative ages of Laurasiatheria and Primates. The Bayesian inference of the relative ages concluded that Primates originated later with

0.99 probability (Supplementary Results and Figs. S22b and S22c). This result emphasises the usefulness of this methodology to assess the relative time of origin of two non-nested clades, with many potential applications in the study of evolutionary radiations and coevolution.

## Discussion

Here, we have formalised and tested the branch length ratio method for the relative timing of evolutionary events based on gene trees (Pittis & Gabaldón, 2016a). A key step in this method is the normalisation of a phylogenetic distance of interest by the median of the branch lengths in a clade, consistently present in all the gene phylogenies. Our results show that, as compared to raw distances, normalised distances serve to better discriminate the timing of evolutionary events. Moreover, we show that these distances, inferred from the distributions of branch lengths in gene tree sets, show a high correlation with molecular clock dating of a species tree based on a concatenated set of single-copy orthologous genes. Finally, we show that our implementation of the approach in a Bayesian framework enables a probabilistic interpretation of the relative timing of events.

The branch length ratio method can use gene family trees that include duplication events. In comparison, molecular dating of a species tree generally relies exclusively on single-copy genes. As a result, the branch length ratio method can exploit a larger amount of available data. We here show that the distribution of normalised distances in collections of gene trees exhibits variability, but consistently has a mode that correlates well to the timing of the studied evolutionary event. We furthermore show that the use of Bayesian inference can account for uncertainty and allow precise estimations of the relative timing of compared events.

Although the purpose of the branch length ratio method is to compare relative timings of events and not provide absolute age estimates, the observed correlation between the relative ages provided by this method and a standard molecular dating approach suggest that this could be possible in some circumstances. A possibility that deserves further investigation. Gene tree branch lengths are primarily determined by the divergence time between sequences and the evolutionary rate of the gene. While evolutionary rates can vary across branches (heterotachy), we demonstrate here that normalizing branch lengths and modelling their distributions provides a measure of relative time, allowing us to detect abnormal branch lengths and discard them.

Regarding this normalisation, we found that the median of the root-to-tip distances within a specific clade conserved across gene trees provides a reliable normalising factor for molecular rate. This is supported by the observation of high correlation between normalized branch lengths and absolute dating based on calibrated molecular clocks. Telford et al. (2014) used the total tree length (the sum of all branch lengths) divided by the number of leaves as a proxy of rate. However, we demonstrate that this measure is strongly affected by heterotachy. Moody et al. (2022) used the mean root-to-tip distance of the minimal ancestor deviation rooted gene tree. While this measure, as the one proposed before, does not account for the asymmetry of the branch length distributions, it offers improved accuracy by considering evolutionary distances from a common ancestral event to the present, albeit one encompassing the entire tree. In our approach, we assume that the distribution of branch lengths from the primates’ most recent common ancestor (MRCA) to the present tips can effectively normalise the gene’s evolutionary rate, similar to the approach taken by Pittis and Gabaldón (2016a) who used paths from the Last Eukaryotic Common Ancestor (LECA) to the present. This type of measure has proven useful for obtaining a relative-to-time estimate by treating branch lengths as random variables with inherent variability.

Recently, Moody et al. (2022) revealed that species trees obtained from concatenated conserved genes result in different estimations for the archaeal-bacterial branch length depending on different gene sets or model parameters. Similarly, Eme et al. (2023) have shown that some species are artifactually close in the species phylogeny when using ribosomal protein sets due to sequence coevolution under similar environmental pressures. Only, when using a functionally broader gene set, they retrieved the currently accepted topology. Thus, the use of larger gene sets, as enabled by the branch length ratio method is likely to alleviate functional and rate biases typical of reduced gene sets.

Here, we modelled evolutionary distances as a gamma-distributed random variable to infer the relative timing of an evolutionary event. Similar to the molecular clock method, we assumed this event was a discrete event, occurring once within a specific timeframe. Consequently, our modelling aimed to retrieve a specific distribution value rather than the entire distribution. Given the predominantly asymmetrical nature of the normalized distributions, we selected the mode, rather than the mean, as a proxy for the event’s timing. As proposed by Martin et al. (2017), we tested the lognormal distribution. Although both lognormal and gamma distributions showed similar behaviours, we believe the latter to be more intuitive as its parametrisation allows us to directly assess the shape and variability of the inferred distribution. Nevertheless, any distribution with positive support and skewness could potentially model the relative dating (once its suitability is proved).

We demonstrate that the use of a Bayesian modelling framework allows assigning probabilities to the comparisons of the timing of two or more events. We show that this approach is robust even when using a limited gene set. The minimum number of trees to obtain a reliable relative time estimate depends on the phylogenetic signal of the used set of genes and genomes and the evolutionary depth of the event. Therefore, each case must be analysed to detect a minimum set of trees to study an event. While the branch length ratio method is susceptible to the presence of high heterotachy leading to skewed normalized branch lengths, such problematic genes can be identified and excluded. Hence, by leveraging comprehensive genome-wide collections of gene trees (phylomes), normalised distances, and a Bayesian modeling framework, the branch length ratio method can obtain reliable relative timing inferences for ancestral events. As shown here, these inferences correlate well with existing dating methodologies, underscoring the validity of the approach. In summary, the branch length ratio method offers a novel and broadly applicable approach, complementary to molecular clock dating, for inferring the relative timing of evolutionary events.

## Acknowledgements

We thank members of the Gabaldón group for insightful discussions. We acknowledge support from the Spanish Ministry of Science and Innovation for grants PID2021-126067NB-I00, CPP2021-008552, PCI2022-135066-2, and PDC2022-133266-I00, cofounded by ERDF “A way of making Europe”; from the Catalan Research Agency (AGAUR) SGR01551; from the European Union’s Horizon 2020 research and innovation programme (ERC-2016-724173); from the Gordon and Betty Moore Foundation (Grant GBMF9742); from the “La Caixa” foundation (Grant LCF/PR/HR21/00737), and from the Instituto de Salud Carlos III (IMPACT Grant IMP/00019 and CIBERINFEC CB21/13/00061-ISCIII-SGEFI/ERDF).

## Author contributions

Toni Gabaldón conceived the idea; Moisès Bernabeu; Carmen Armero and Toni Gabaldón designed the methodology; Moisès Bernabeu programmed and performed the computations; Moisès Bernabeu, Carmen Armero and Toni Gabaldón analysed the data; Moisès Bernabeu wrote the first draft of the manuscript. All the authors contributed substantially to the draft and gave final approval for publication.

## Data and code availability

The raw output data is stored in the Zenodo repository https://zenodo.org/records/14608060?preview=1&token=eyJhbGciOiJIUzUxMiJ9.eyJpZCI6IjM2MTEyMDYxLWRkY2ItNDJhNi05YThlLTQyYTYwODI5Y2VkYyIsImRhdGEiOnt9LCJyYW5kb20iOiI2ODNmYzBlYjU2MGM2ZmFlMzVmNDY5YjdlMWJiNmQ0YSJ9.memhSxMRaqJNNRYZUVOj9vgue_H0FdeEU1uXIIc7zevxHqKDo27y-CoCqV0Pl5CjtJM2UKetPWhz47dU9TZ21g. The code used for the distances calculation and the posterior distributions calculation is available in the attached compressed file.

