## Supplementary text and figures for "Probabilistic modelling improves relative dating from gene phylogenies"

June 13, 2025

**3 Supplementary tables** **6**

**4 Supplementary figures** **7**

**5 Datasets** **22**

### 1 Supplementary methods

#### 1.1 Posterior distribution of the difference between modes

We computed the posterior distribution of the difference between modes by applying the posterior distribution of the subtraction of the modes for all the combinations of parameters and the probability associated to them.

$$\begin{aligned}\pi(\text{mode}_1 - \text{mode}_2 \mid \mathcal{D}) = \pi\left(\frac{\alpha_1 - 1}{\beta_1} - \frac{\alpha_2 - 1}{\beta_2} \mid \mathcal{D}\right) P\left(\alpha_1 \geq 1, \alpha_2 \geq 1 \mid \mathcal{D}\right) + \\ + \pi\left(\frac{\alpha_1 - 1}{\beta_1} \mid \mathcal{D}\right) P\left(\alpha_1 \geq 1, 0 < \alpha_2 < 1 \mid \mathcal{D}\right) + \\ + \pi\left(-\frac{\alpha_2 - 1}{\beta_2} \mid \mathcal{D}\right) P\left(0 < \alpha_1 < 1, \alpha_2 \geq 1 \mid \mathcal{D}\right)\end{aligned} \tag{1}$$

#### 1.2 GO term enrichment analysis

We annotated the Gene Ontology (GO) terms of each human seed protein using InterProScan v5.39-77.0 (Jones et al., 2014) and performed a GO enrichment analysis using the R package topGO (Alexa and Rahnenfuhrer, 2023) with the method "weight01" and the Fisher's exact test. For this analyses, we set a significance threshold of 0.05 to consider an enriched category.

#### 1.3 Testing on simulated gene tree data

To test the branch length ratio method in more controlled conditions and assess the impact of different duplication and loss parameters, we simulated trees under two different loss and duplication rates using as a reference the dated species tree. The simulations were carried out using SaGePhy (Kundu and Bansal (2019)) with default parameters of the birth-death process but different loss and duplication rates. To confer a degree of variability to the loss and duplication parameters, we determined these values for each simulation using uniform distributions. We generated two sets of 5,000 trees using the SaGePhy GuestTreeGen program. The first set was simulated under high loss and duplication rates ($U(0, 2)$). The second set of simulations was inferred using low loss and duplication rates ($U(0, 0.5)$) and lower rate heterogeneity. This procedure generates an ultrametric tree. Once the trees were simulated, we rescaled their branch lengths using two gamma-modelled rate heterogeneity specifications with SaGePhy BranchRelaxer. We used a $U(1, 4)$ and $U(0, 2)$ for the two parameters of the gamma distribution in the high and low loss / duplication sets, respectively. With this procedure, we obtained simulated gene trees with scaled branch lengths, resembling those of our real data. To assign a seed sequence to each tree, we firstly identified any primates group in the tree, randomly chose one of them and got the human tip as seed. In case of more than one human tips, we randomly picked one. For the simulated trees with a proper primates' node, we computed the tip-to-internode distances.

### 2 Supplementary results

#### 2.1 The behaviour of the normalisation factor

Regarding the behaviour of the normalising factor, we performed a set of correlations between the normalising factor and other definitions for the gene tree rate in the literature. These rate definitions are divided into two tree measures: (i) the total tree length, the sum of all branch lengths, and (ii) the root-to-tip distances. We assessed the symmetry of both measures by getting the correlation between the mean and the mode and the skewness distribution (Supplementary Figs. 12 and 13, respectively). We see that the root-to-tip distance distributions are more symmetrical than the branch lengths, which means that root-to-tip distances are less sensitive to heterotachy. Furthermore, conceptually, they are a better proxy of time as it retrieves the relative measure of a given evolutionary process to the current time (event-to-tip), as the previous formalisation of the branch length ratio method stands for (Susko et al., 2021).

To compare the normalisation factor studied in this paper with the rate defined as the tree length divided by the number of leaves, we performed the correlation between both measures. For the entire tree (Supplementary Fig. 14a), the relationship between the tree length divided by the leaves number and the normalising factor is sparse. In contrast, both variables become more related when this ratio is considered for the primates subtree. As a definition of evolutionary rate for a gene tree, we think it is quite reductionist, given that, in fact, it is the scaled mean branch length (Supplementary Fig 17). It does not account for evolutionary events timing. Still, the singular protein change in each branch could be biased by underlying specific evolutionary processes or inference artifacts such as misestimation of the branch length due to model selection. Furthermore, the mean is extremely sensitive to outliers, and as we have proven, branch length distributions are mostly skewed.

#### 2.2 Impact of mixture models in normalised branch lengths

Branch lengths are sensitive to the model used in phylogenetic inference. Mixture models account for equilibrium frequencies (CXX family, e.g. LG+C60+G4), and exchangeabilities (LG4X or LG4M) differences across alignment positions. However, they are far more computationally expensive. To test such mixture models in our alignments and assess its effect in the computed normalised distances, we randomly selected 1,000 alignments and ran two different IQ-TREE commands. The first command allowed for model selection among the previously used models (see Methods) and LG+C60+G4, LG4X and LG4M:

```
iqtree -s {aln} -pre {prefix} -madd LG4X,LG4M,LG+C60+G4
-mset DCmut,JTTDCMut,LG,WAG,VT -mfreq F -mrate E,G,R,I+R,G+R
-m MFP -nt 4
```

In the second IQ-TREE configuration, we fixed the model to LG+C60+G4 for all the alignments:

```
iqtree -s {aln} -pre {prefix} -m LG+C60+G4 -mfreq F -nt 4
```

When allowing model selection using ModelFinder, LG+C60+G4 is selected just in 7 out of the 1,000 alignments, LG4M in 28, and LG4X in 29. For the rest of the alignments (93.6%) the model used in the original run (without mixture models in the models set) was selected as the best model. After two days of computing using 8 threads running 14 simultaneous trees, just 101 trees finished. The median runtime of the best-fit model was 380.57 seconds, whereas for the forced LG+C60+G4 model was 68,199 seconds, 180 times slower. This computational effort does not help to improve, nor change, the general trends regarding branch lengths, as we observe a strong correlation between the comparable branch lengths using CXX models and without using them (Supplementary Figure 18).

#### 2.3 Use of alternative normalising clades

To test whether the use of a normalising clade nested inside the events of interest was driving the estimate of the age and assess the impact of the use of alternative normalising clades, we repeated the analysis using two different normalising clades, not nested inside the events of interest: Laurasiatheria and Rodentia. Despite some dispersion, the distances normalised with primates show a significant positive correlation with both Laurasiatheria ($\rho = 0.49$, $p \leq 2.2e - 16$) and Rodentia ($\rho = 0.43$, $p \leq 2.2e - 16$) (Supplementary Figs. 20a and b). The positive correlation indicates that any of these normalising clades, when found in the phylogenies, result in measures that can be related to time, as they correlate to those analysed in the current study. Therefore, the normalisation based on a clade common to all the analysed gene trees is robust regardless of the chosen reference clade. More importantly, the distributions of normalised distances using any of the normalising clades were highly similar and reflected the same ordering of events (Supplementary Figs. 20c and d).

##### 2.4 Testing on simulated gene tree data

We simulated two sets of 5,000 trees using two different specifications for the duplication and loss parameters and gamma rate heterogeneity. The first set had high values for all the parameters, and the second set had low values for all the parameters.

In the simulated trees, we observe differences in the number of simulated trees that have a proper Primates node. Higher duplication and loss rates result in a low fraction of trees with the Primates node (1,422 out of 5,000 simulated trees). More losses and duplications may result in topological incongruences with the species tree that hinder the identification of a Primates node. In the case of the lower rates simulations, we observe a larger fraction of trees congruent with the species tree (3,588 out of 5,000 simulated trees).

As observed in real data, the simulated trees provide tip-to-internode raw distances distributions that are harder to distinguish for each event than normalised distances distributions. The change of rates of transfer, loss and gamma rate heterogeneity cause changes in the raw distances magnitude. Lower values for the rates provide shorter raw distances, while higher rates provide longer raw distances.

The effect of normalisation resembles that in trees reconstructed with real sequences. The distributions are narrower and they are more separated. However, simulated trees show closer distributions between events than real data trees. Simulations also show that older events have more uncertainty, as the distribution is more dispersed. The simulated trees using lower rates provide wider distributions when compared to trees simulated with higher rates.

Overall, simulations support that normalisation provides distributions for the events that follow the species tree ordering. However, we observe that real data distributions resolve better the differences between the studied events. This may reflect that simulations are not realistic enough and more dependent on the simulation parameters, such as the branch lengths of the starting species.

##### 2.5 Comparing relative ages of non-nested clades

To compare whether the described method allows us to compare two non-nested events and elucidate the probability of one occurring before or after the other, we calculated the distances from the most recent common ancestor of Primates and Laurasiatheria (Placentalia) to each of these clades. This distance provides us a framework to assess whether Laurasiatheria predated Primates or not, as the reference clade would be their ancestor. In contrast to distances from the seed to the event of interest, here the distance interpretation is the opposite. Longer normalised distances for an event would mean that the clade originated later, whereas shorter distances would show an earlier origin of the clade. Therefore, using these MRCA-to-internode distances, we can compare the origin of two non-nested clades.

The empirical distributions for the normalised distances from the MRCA to Laurasiatheria and Primates have longer tails than the tip-to-internode distances (Supplementary Figure 21a). These long tails hinder the possibility to assess the sorting. As the distribution showed a first peak (vertical line for lower values in Supplementary Figure 21a), we iterated through different thresholds of maximum normalised distance based on the quantile of normalised distances. When removing those normalised distances longer than the 35th quantile, we distinguished between both distributions (Supplementary Figure 21b).

Although the fraction of trees that can be used for these analyses is much lower than for the tip-to-tip and tip-to-internode distances, we can better differentiate between both distances when focusing on the peak of the distribution. The normalised filtered distances, again, agree with the molecular clock ages, concluding that Laurasiatheria clade origin (83.3 – 88.0 Mya) predates the origin of Primates (71.4 – 77.5 Mya).

Doing a similar procedure as the tip-to-internode distance, we calculated the distances from human to Primates and from cat to Laurasiatheria using Placentalia as the normalising clade, which is common to both events. In this case, longer distances would mean an older event, whereas shorter distances would mean a more recent event. We observe that Laurasiatheria presents longer branch lengths than Primates (Supplementary Figure 22a). Agreeing with both the MRCA-to-internode distances and also the species tree.

We performed the Bayesian analysis on the tip-to-internode of different clades, and we could compute the posterior distributions of the mode for Laurasiatheria and Primates, and assess the sorting of these clades (Supplementary Figure 22b and c). This analisis resulted in a older age estimate for the Laurasiatheria node, whereas Primates showed shorter normalised branch lengths. When assessing the hypothesis of primates being older than laurasiatherians, the probability is 0.0004, agreeing with the hypothesis suggested by the molecular clock tree. These results show that the Bayesian implementation of the branch length ratio method also helps resolve evolutionary hypotheses with non-nested clades. Being able to compare these hypotheses accounting for more genes, may provide insights into several competing clades such as fungi and plants, or even in coevolution studies. In the latter, one could compare whether two coevolving clades have similar diversification timelines.

#### 3 Supplementary tables

These supplementary tables are in the attached spreadsheet.

Supplementary Table 1: Taxon sampling lineage, used Uniprot code and NCBI TaxID.

Supplementary Table 2: Summary, convergence statistics and autocorrelation values for the seed-to-event distances Gamma and Normal inferences posterior MCMC samples. SD: standard deviation, R: Gelman-Rubin's statistic, ESS: effective sample size, a: alpha, b: beta, m: mean, mo: mode and v: variance.

Supplementary Table 3: Summary, convergence statistics and autocorrelation values for the seed-to-species distances Gamma and Normal inferences posterior MCMC samples. SD: standard deviation, R: Gelman-Rubin's statistic, ESS: effective sample size, a: alpha, b: beta, m: mean, mo: mode and v: variance.

Supplementary Table 4: Summary, convergence statistics and autocorrelation values for the seed-to-lineage node distances Gamma and Normal inferences posterior MCMC samples. SD: standard deviation, R: Gelman-Rubin's statistic, ESS: effective sample size, a: alpha, b: beta, m: mean, mo: mode and v: variance.

Supplementary Table 5: Variation percentage between the full tree set and the 10% of trees set for the standard deviation (SD), the minimum, maximum and mean values for the posterior distribution of the events.

Supplementary Table 6: Variation percentage between the full tree set and the 10% of trees set for the standard deviation (SD), the minimum, maximum and mean values for the posterior distribution of the comparison of events.

### 4 Supplementary figures

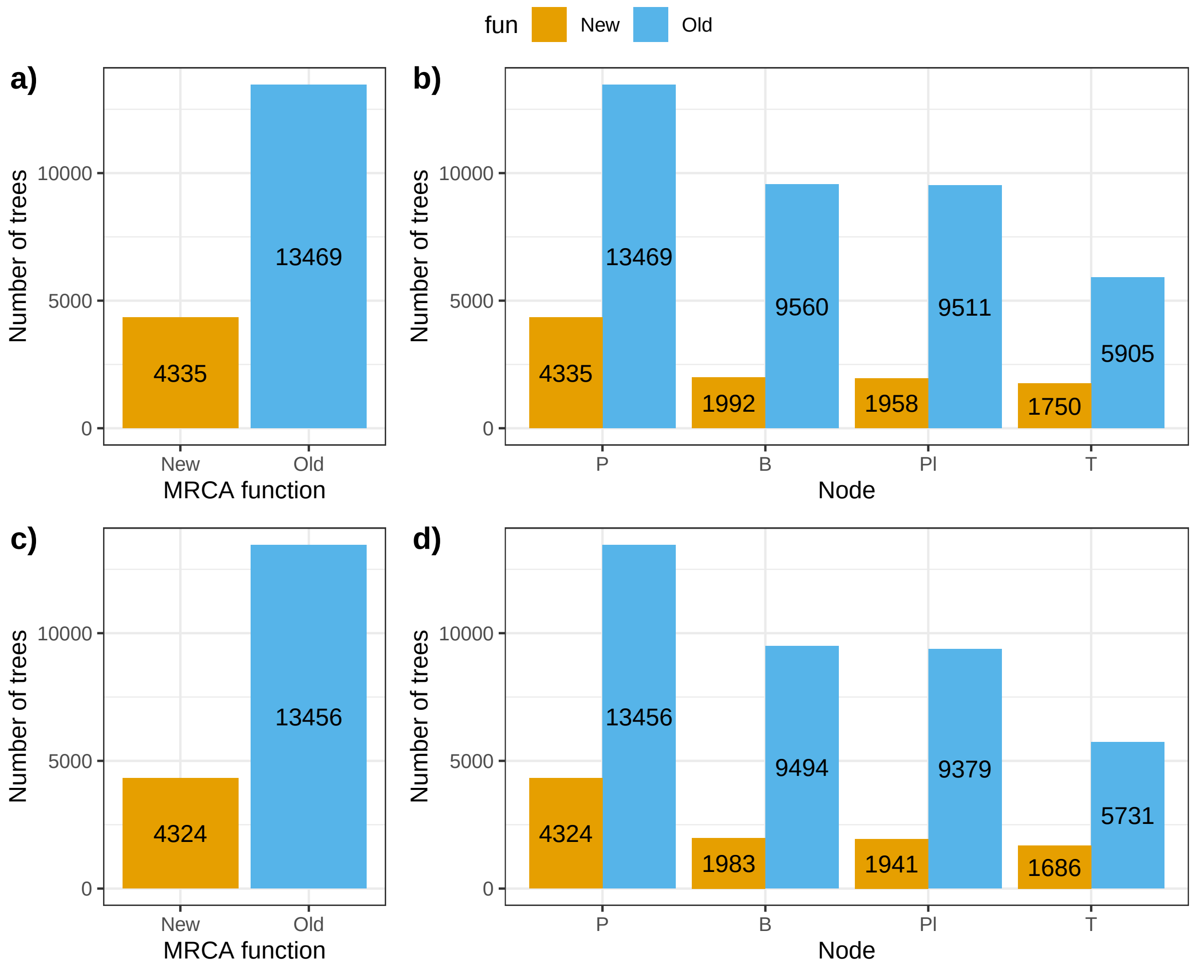

Supplementary Fig. 1: The number of trees per function before (a, b) and after filtering (c, d). a) The number of trees by MRCA definition function and b) the number of trees containing the x-axis node without filtering. a) The number of trees by function and b) the number of trees containing the x-axis node above the quantile 0.99 of the normalised distances. The new function only defines a node as the event's node if it matches the species tree event's first split species composition, the old one is a classical MRCA function which assigns the event given a set of species belonging to such event without taking into account the species tree topology.

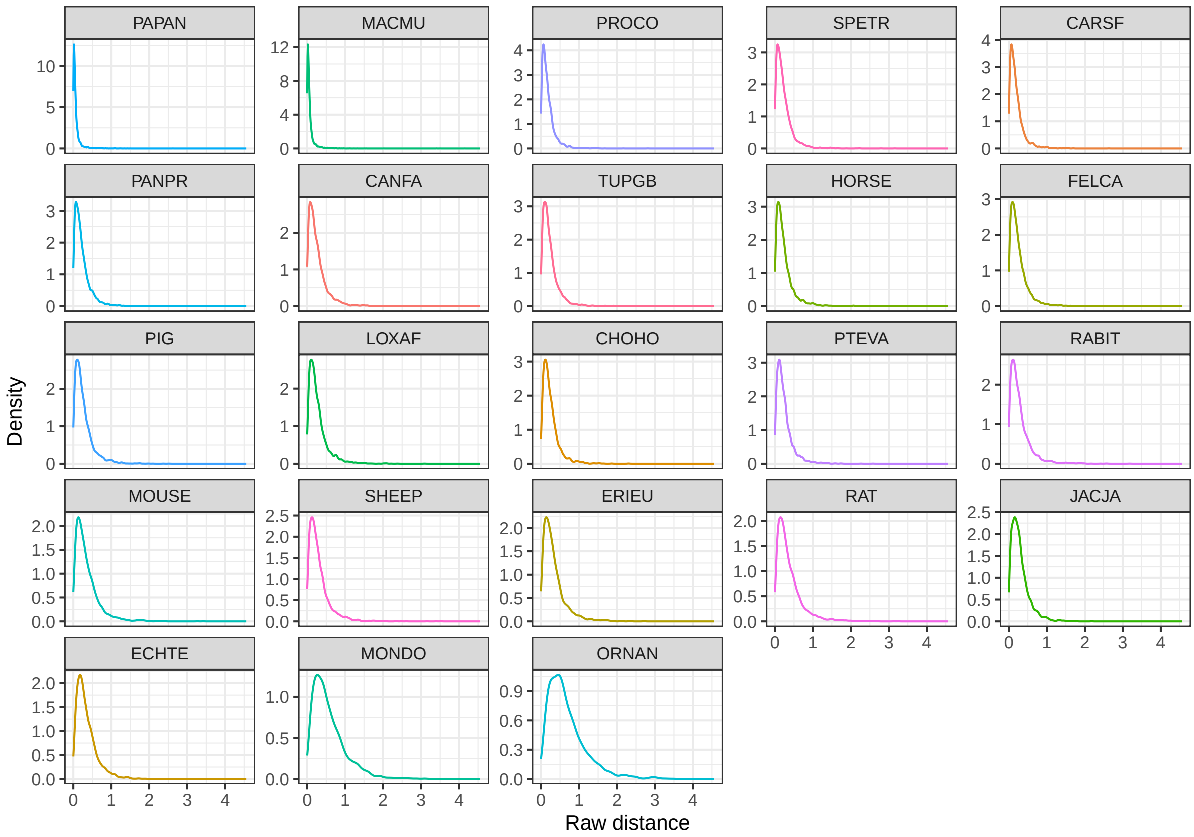

Supplementary Fig. 2: Raw tip-to-tip distances distributions. Each box shows the distribution of the distances from the human seed to the species indicated. These distributions come from distances between orthologous and co-orthologous sequences (see methods). The distributions are sorted by mode.

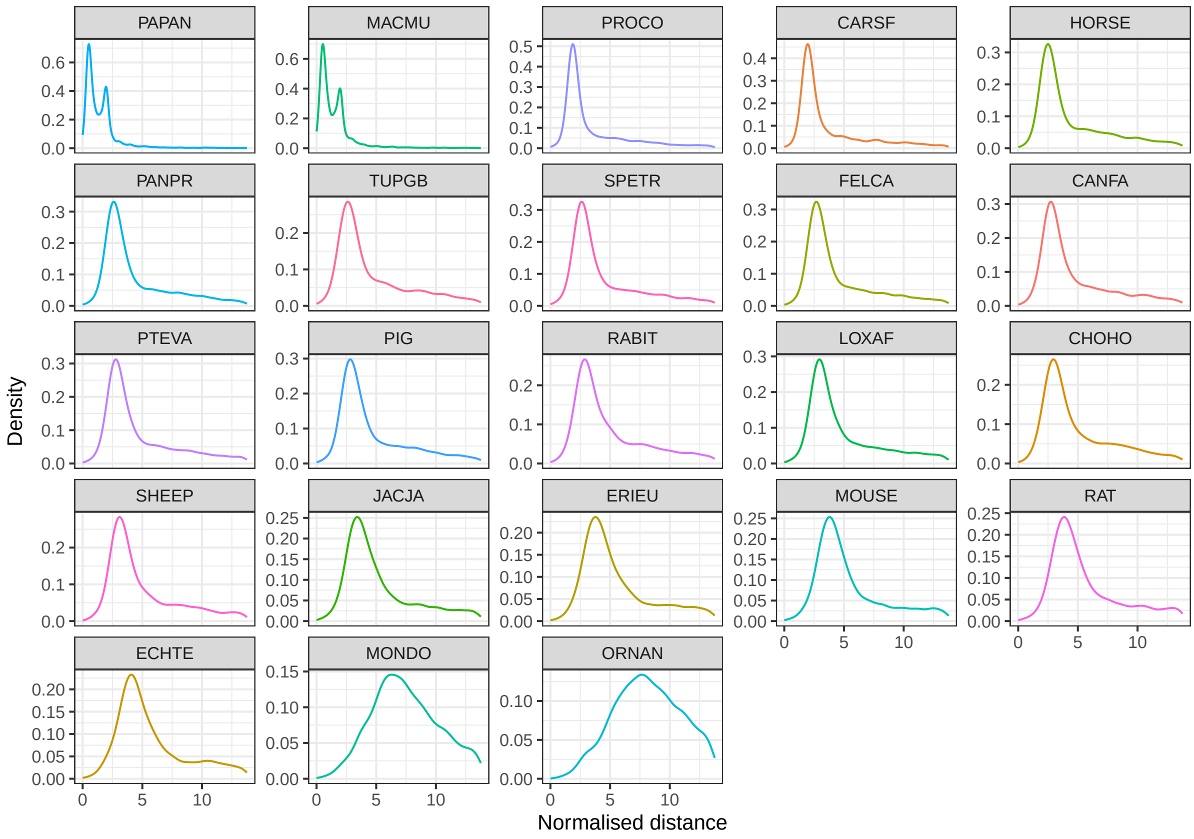

Supplementary Fig. 3: Normalised tip-to-tip distance distributions. Each box shows the distribution of the distances from the human seed to the species indicated normalised by dividing the raw distances by the branch lengths median in the Primates clade. These distributions come from distances between orthologous and co-orthologous sequences (see methods). The distributions are sorted by mode.

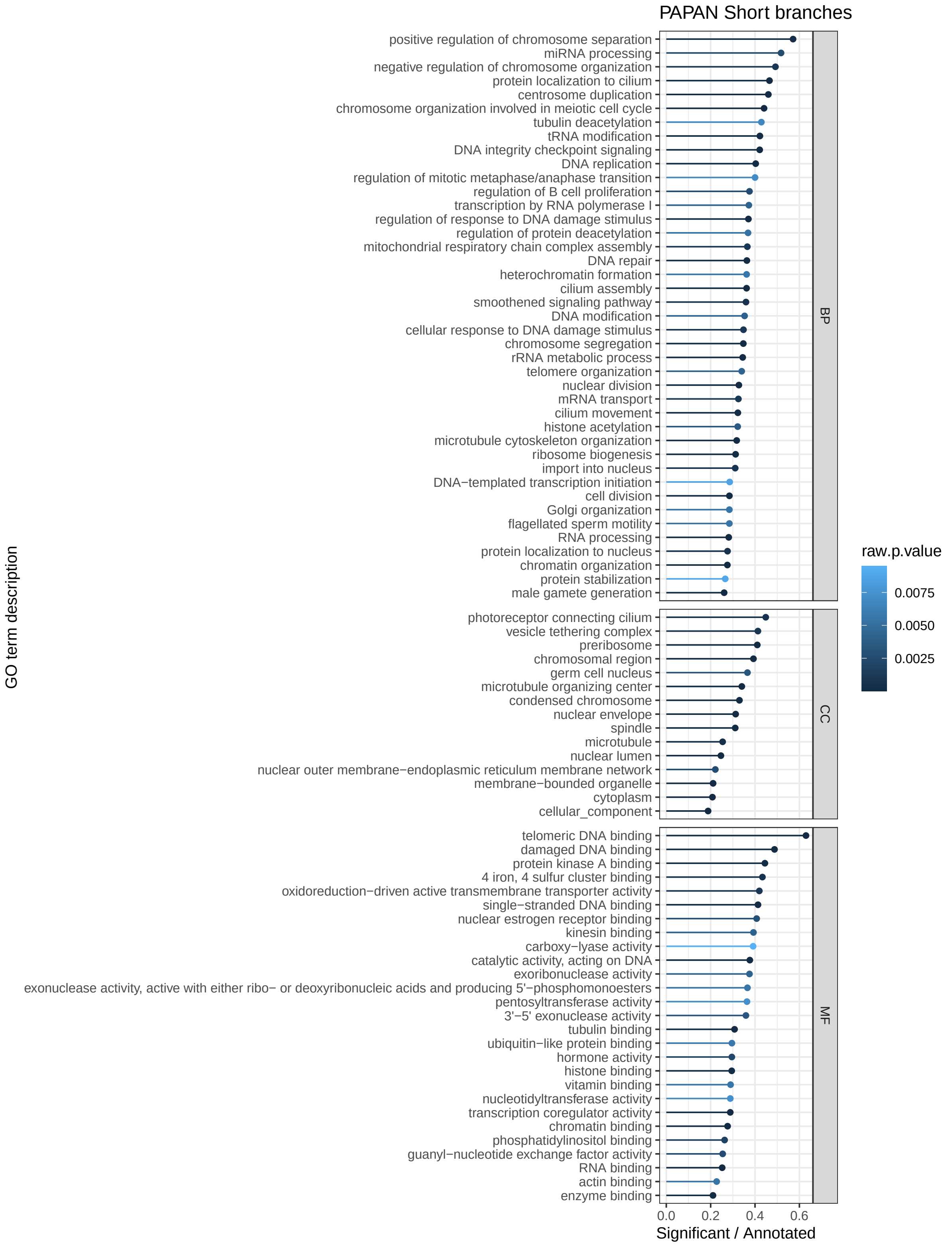

Supplementary Fig. 4: *Papio anubis* first peak GO enrichment. GO enrichment for the genes in the first peak (shorter normalised distances) of the bimodal distribution.

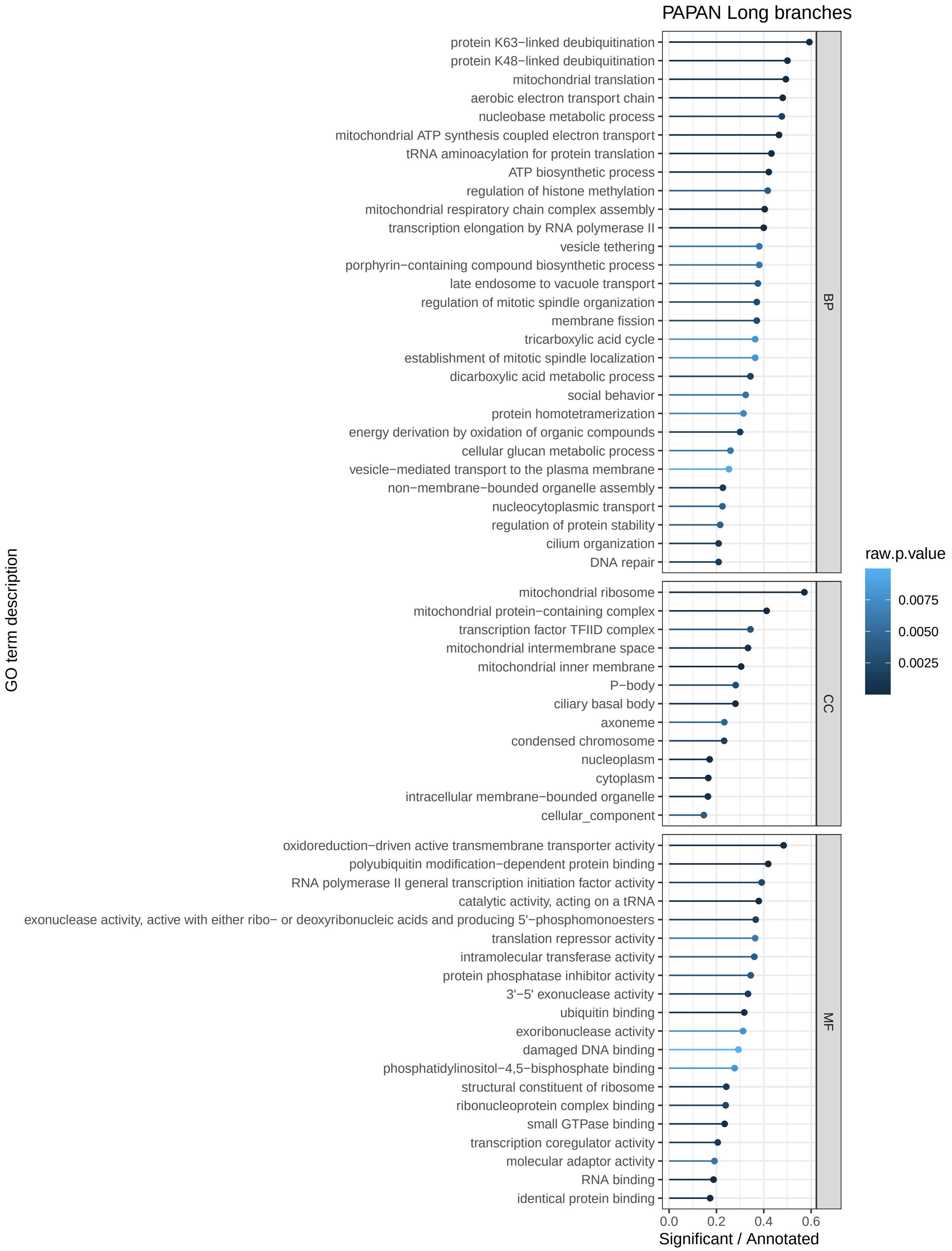

Supplementary Fig. 5: Papio anubis second peak GO enrichment. GO enrichment for the genes in the second peak (longer normalised distances) of the bimodal distribution.

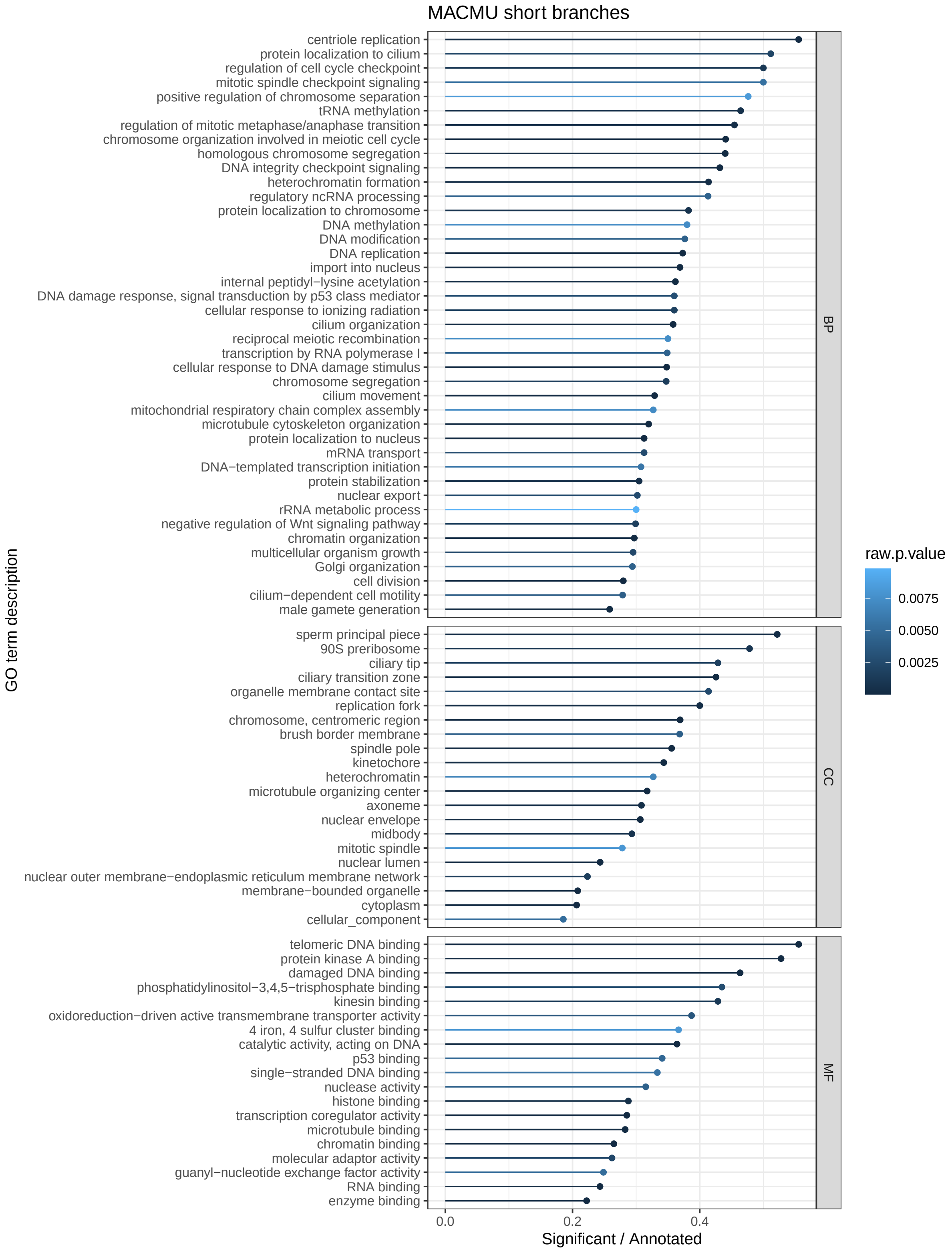

Supplementary Fig. 6: *Macaca mulatta* first peak GO enrichment. GO enrichment for the genes in the first peak (shorter normalised distances) of the bimodal distribution.

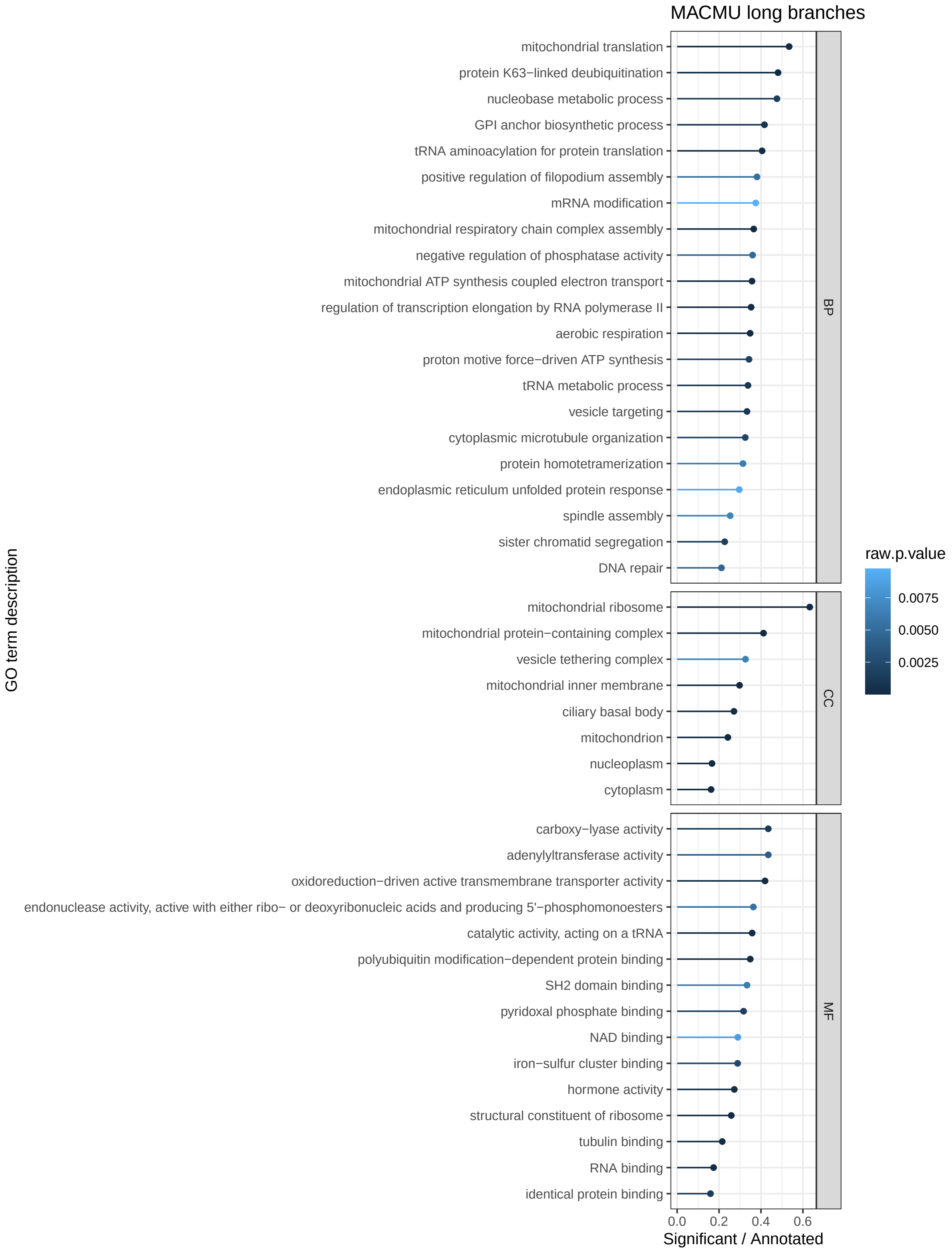

Supplementary Fig. 7: *Macaca mulatta* second peak GO enrichment. GO enrichment for the genes in the second peak (longer normalised distances) of the bimodal distribution.

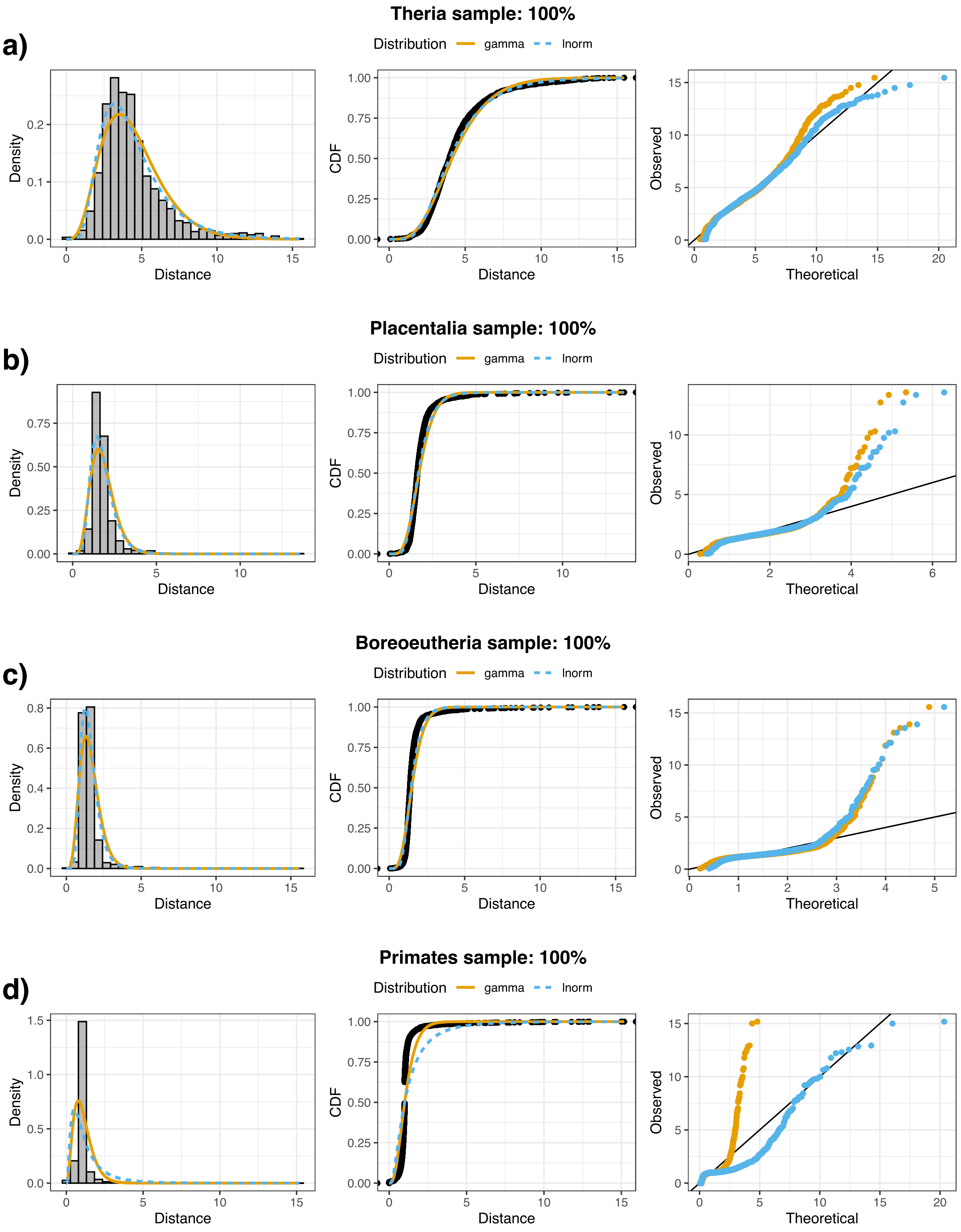

Supplementary Fig. 8: Inferred distributions and their fitting to the observed distances histograms. Seed to a) Primates, b) Boroeutheria, c) Placentalia and d) Theria event normalised distances histograms in the left, in the middle the quantile-quantile plot and in the right the CDF plot. The orange lines and points identify the gamma distribution, and the blue dashed lines and points identify the lognormal distribution.

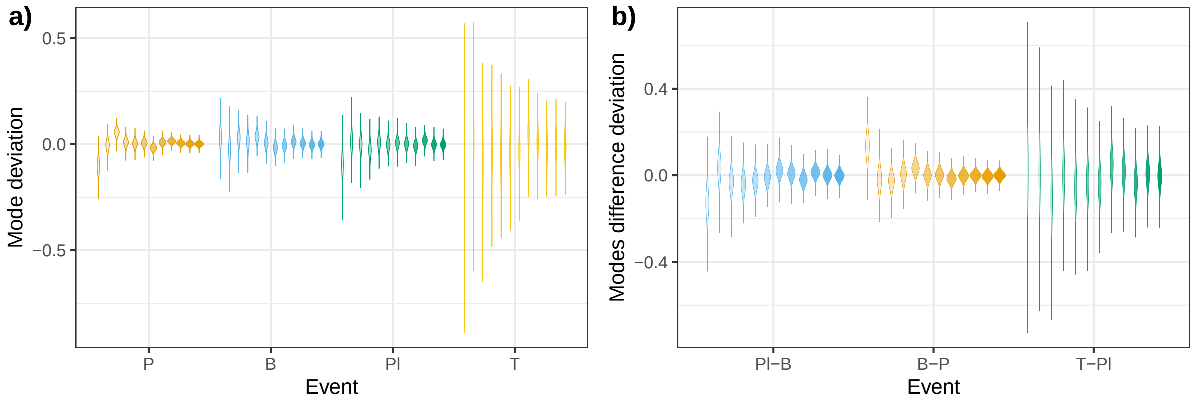

Supplementary Fig. 9: Subsampling divergence with the full tree set. Violin plots a) for the posterior distributions of the mode of each event and subsample centred in the whole tree set as a measure of the deviation from the inferred mode, and b) for the posterior distributions of the mode difference between contiguous events centered to the mode of the full tree set. Subsampling range from 10% of the total number of trees to 100%, increasing by 10% each, we also included 15% and 25% of the trees. The more opacity of the violin, the higher percentage of trees used. Each colour shows the event and comparison according to the x-axis.

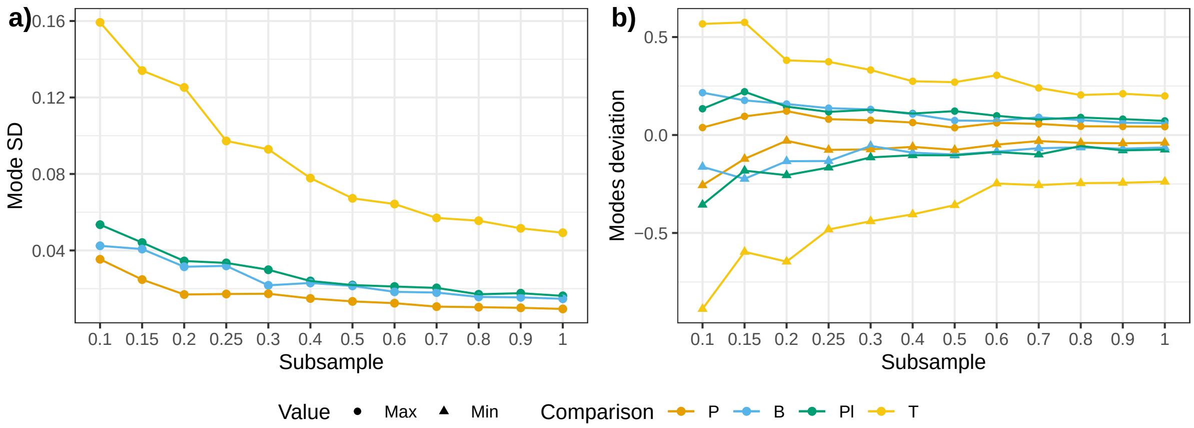

Supplementary Fig. 10: Events dispersion through subsamples. a) Evolution of the posterior distribution standard deviation for each event. b) Evolution of the maximum (point) and the minimum (triangle) values of the posterior distribution of the event's mode centred in the full tree set (0 means the inferred mode with all the trees). Each colour shows the event according to the legend.

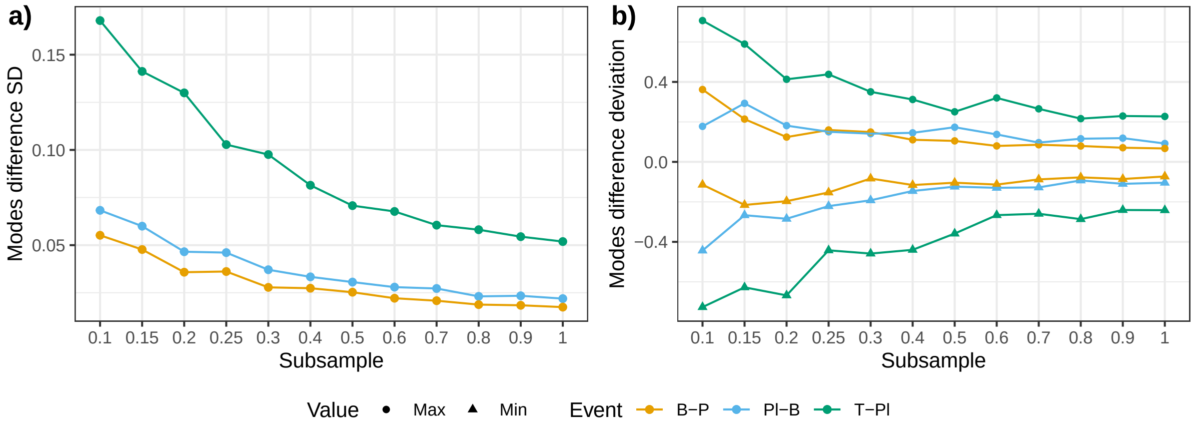

Supplementary Fig. 11: Comparison of events dispersion through subsamples. a) Evolution of the posterior distribution standard deviation for each subsample. b) Evolution of the maximum (point) and the minimum (triangle) values of the posterior distribution of the modes comparison centred in the full tree set (0 means the inferred modes' comparison with all the trees). Each colour shows the comparison according to the legend.

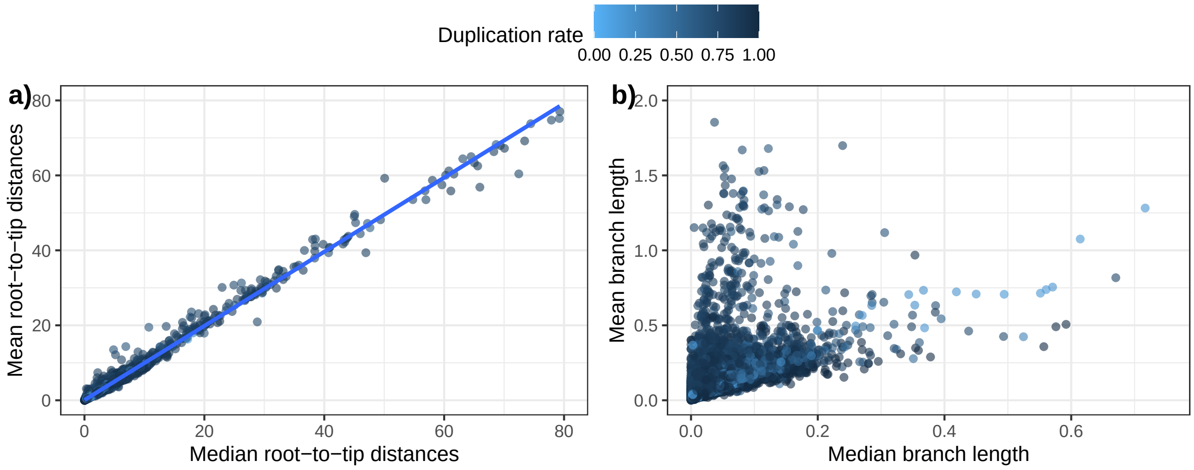

Supplementary Fig. 12: Symmetry assessment of the distribution of some tree measures. Correlation between the median and the mean of the a) root-to-tip distances and b) branch lengths. Colour intensity shows the duplication rate, the darker, the more duplicated.

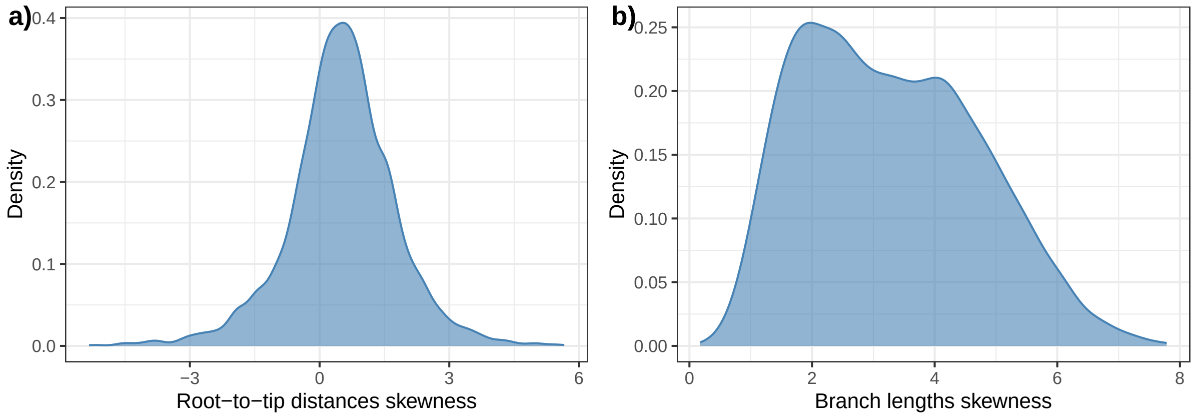

Supplementary Fig. 13: Skewness distribution. a) For the root-to-tip distances and b) for the branch lengths across the phylome whole trees.

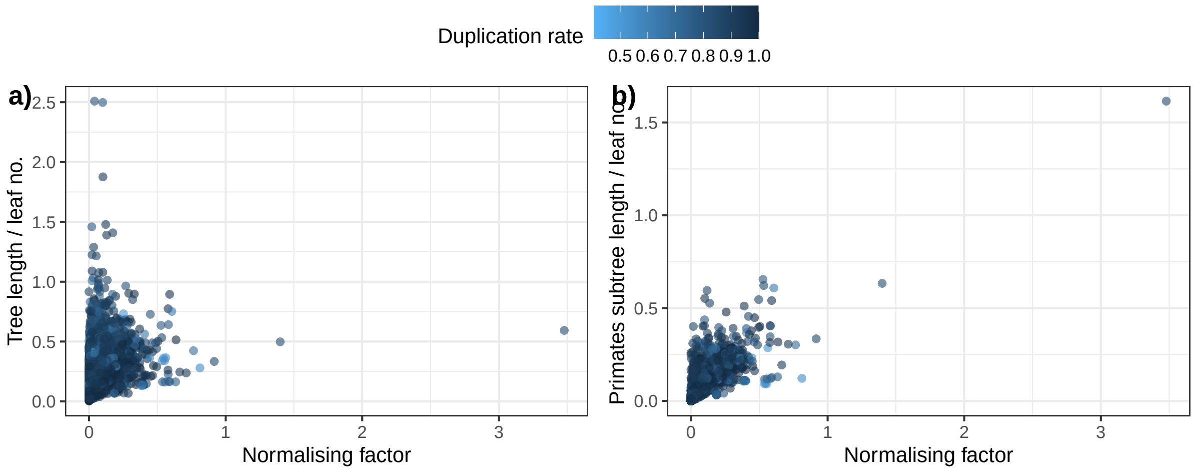

Supplementary Fig. 14: Normalising group correlations. Normalising factor (median of the root-to-tip distances for the primates clade) correlation with a) the tree length divided by the number of leafs and b) the primates subtree length divided by the number of leafs. Colour intensity shows the duplication rate, the darker, the more duplicated.

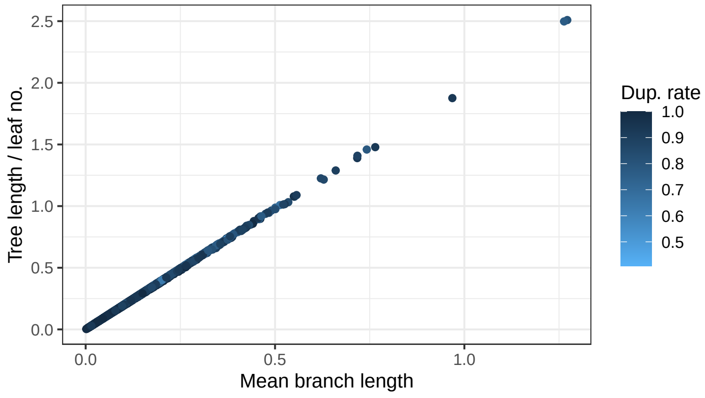

Supplementary Fig. 15: Mean branch length correlation with the tree length divided by the leaves number. Colour intensity shows the duplication rate, the darker, the more duplicated.

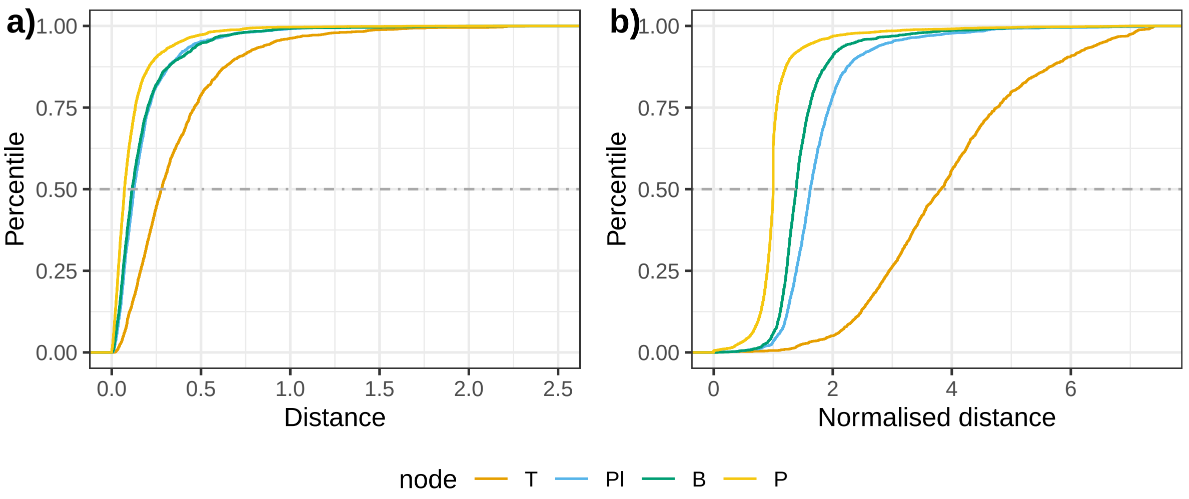

Supplementary Fig. 16: Empirical cumulative distribution functions (ECDFs) for the a) raw and b) normalised distances.

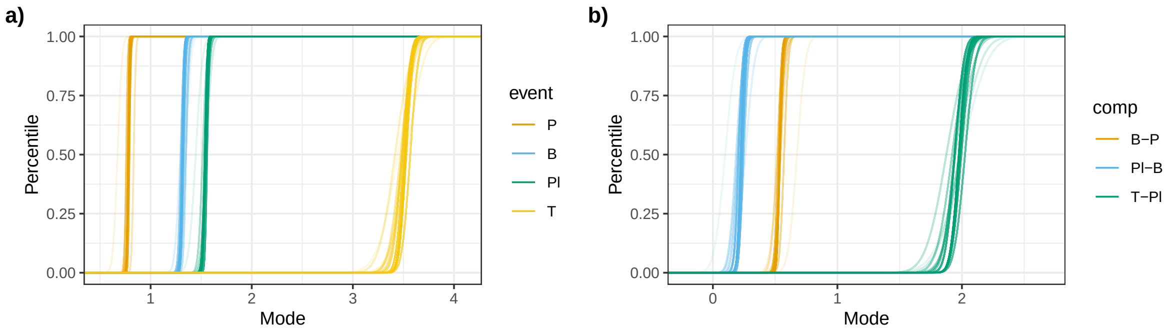

Supplementary Fig. 17: Empirical cumulative distribution functions (ECDFs) for the posterior distributions of the a) modes of each event, and b) comparison between modes.

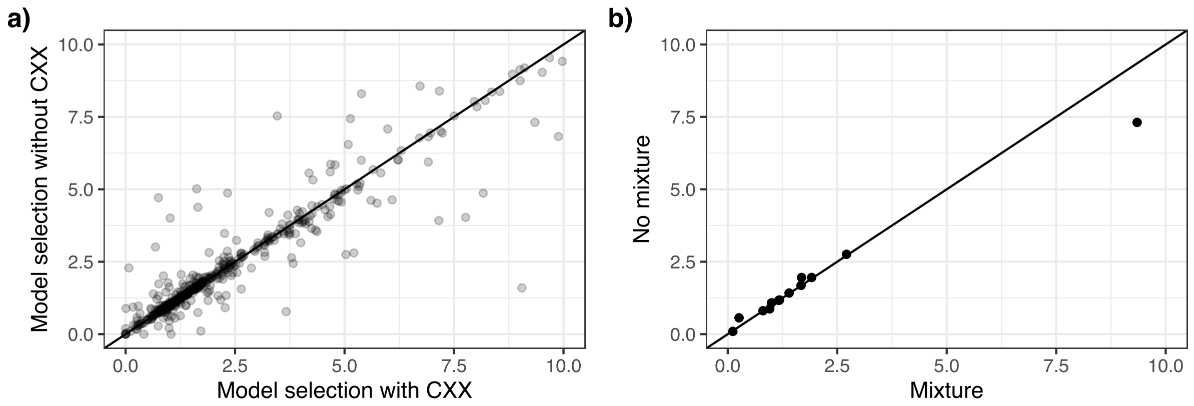

Supplementary Fig. 18: Correlation of normalised distances between mixture and non-mixture models. a) Correlations between equivalent distances in the same tree inferred under model selection including mixture models (LG4X, LG4M and LG+C60+G4) and empirical equilibrium frequencies (+F) and without including these models and parameters. b) Correlation of comparable distances present in trees that changed from no mixture models to mixture models.

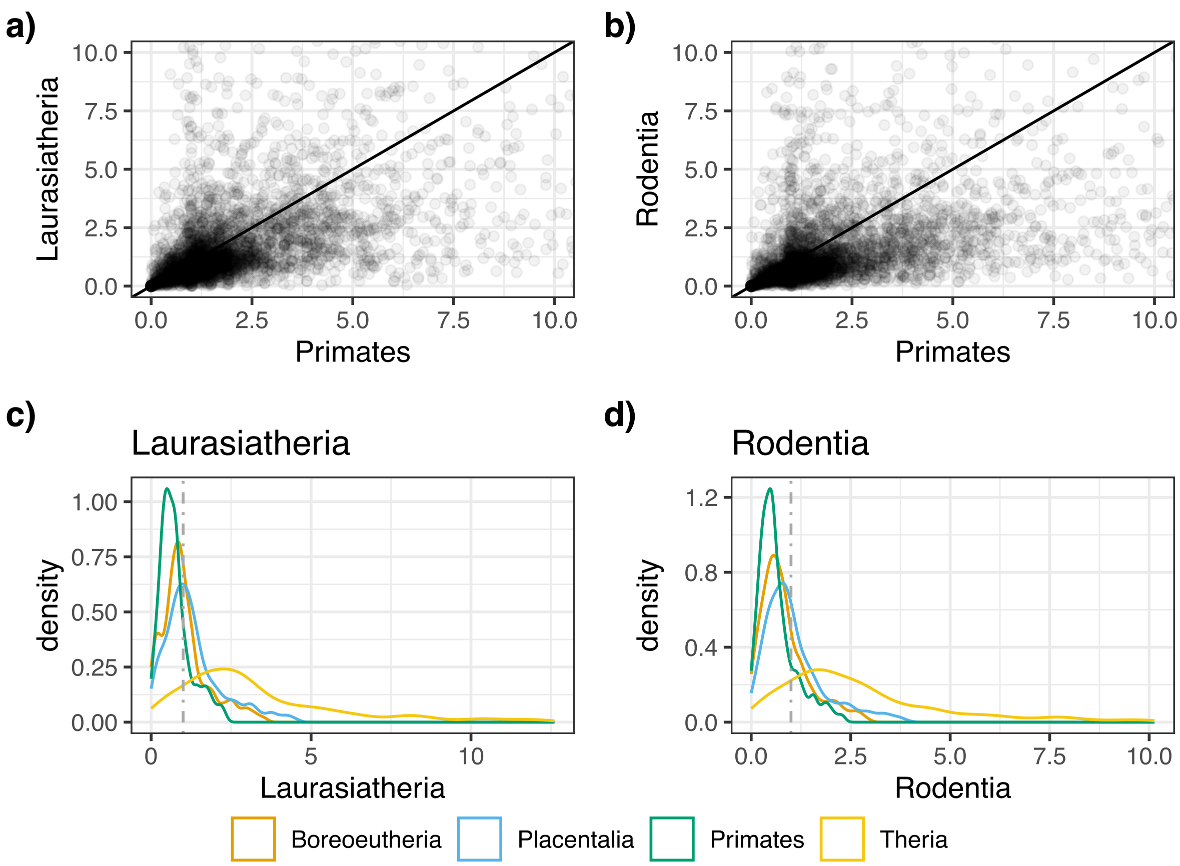

Supplementary Fig. 19: Impact of different normalising groups in the normalised distances. a) Correlation between the distances from human to internal nodes of interest using Primates or Laurasiatheria as normalising clade. b) Correlation between the distances from human to internal nodes of interest using Primates or Rodentia as normalising clades c) Distributions of the normalised distances from human to the events of interest using Laurasiatheria as the reference clade. d) Distributions of the normalised distances from human to the events of interest using Rodentia as the reference clade. Vertical lines in c) and d) indicate a normalised distance value of 1.

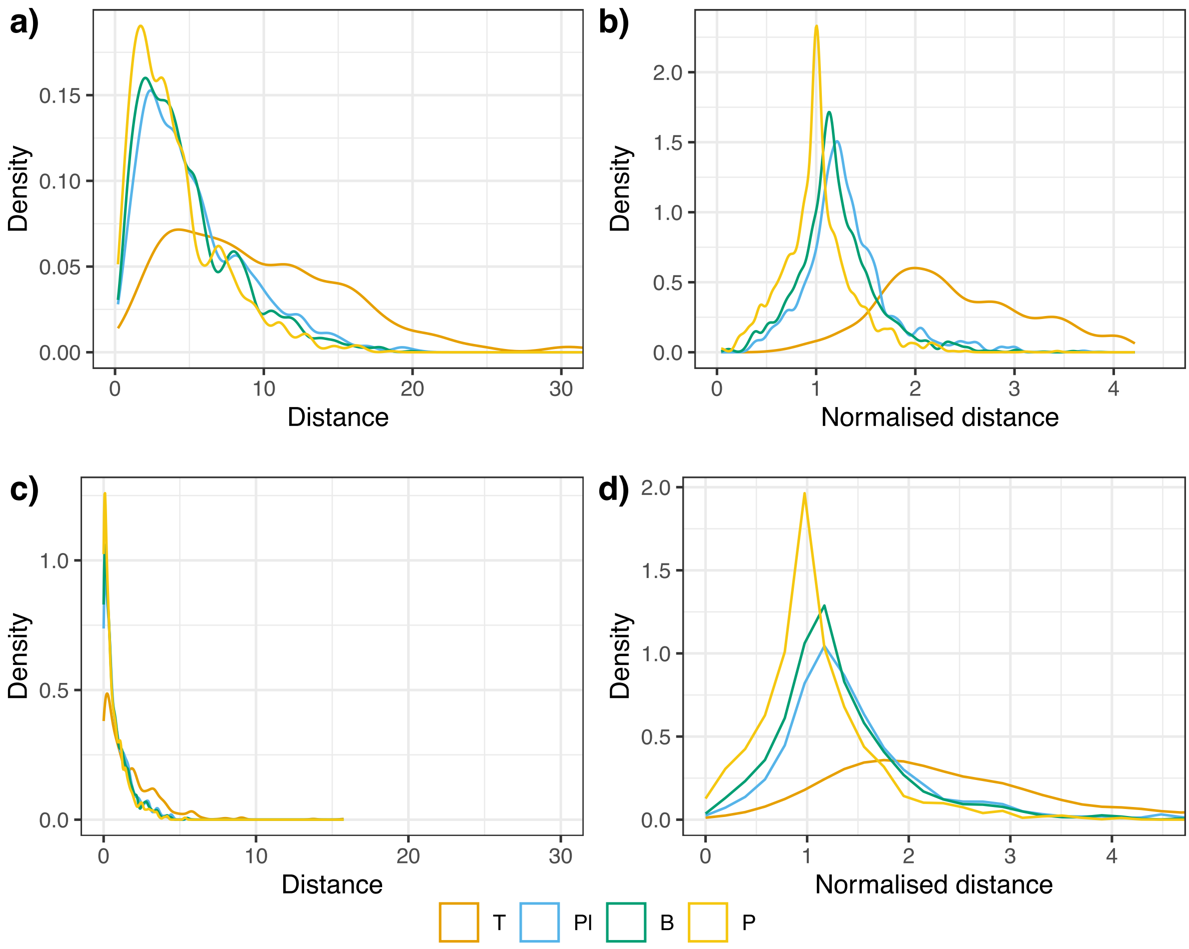

Supplementary Fig. 20: Normalisation effect in simulated data. a) Raw and b) normalised distances in a fast-evolving and high loss and duplication rates set of simulated trees. c) Raw and d) normalised distances in a slowly-evolving and low loss and duplication rates set of simulated trees.

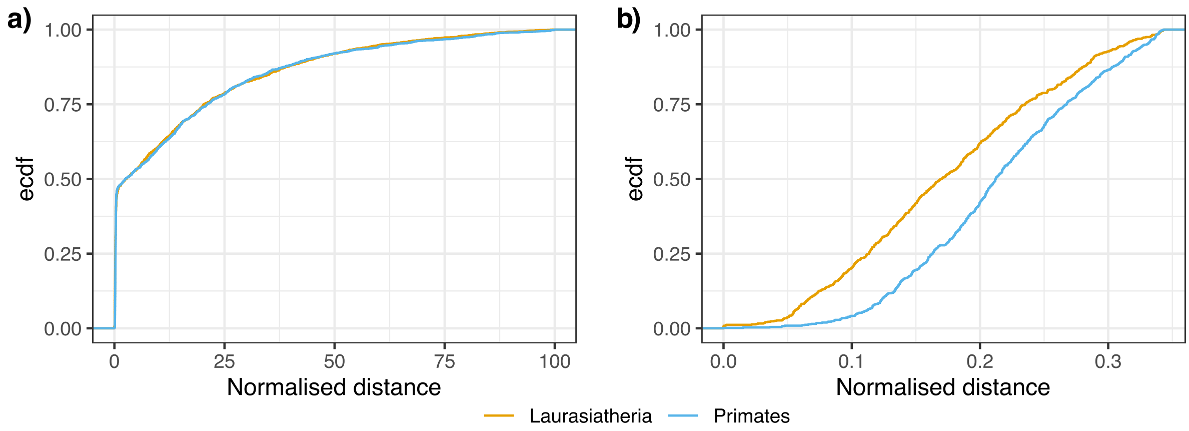

Supplementary Fig. 21: Empirical cumulative distribution functions of the MRCA-to-internode normalised distances for two competing nodes, Laurasiatheria and Primates. a) ECDFs for normalised distances, all the trees with both distances are included, except those with a normalised distance longer than the 90th percentile, as for the tip-to-event distances. b) ECDFs for the normalised distances of those trees with a distance lower than the 35th percentile.

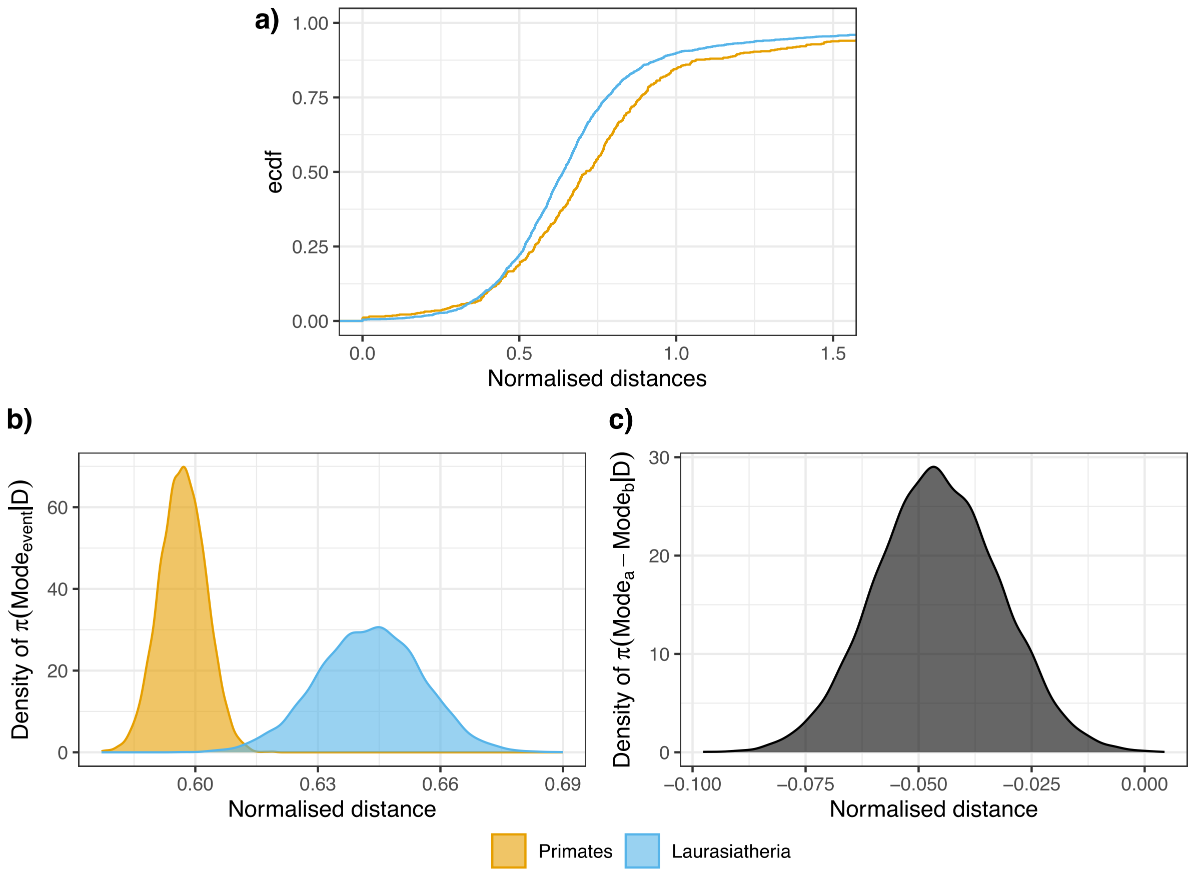

Supplementary Fig. 22: Non-nested clades comparison. a) Empirical cumulative distributions for the normalised tip-to-internode distances from cat to Laurasiatheria and human Primates node. The distances are normalised using the Placentalia node. b) Posterior distributions of the mode of each event. c) Posterior distribution of the comparison between modes.

### 5 Datasets

Stored at the Zenodo Repository (link).

- mammal_trees.zip: compressed trees form the Human phylome. It can also be downloaded from the PhylomeDB website with the code: 0593.
- distances.zip: raw output of top-to-internode (events and lineage), tip-to-tip distances R data frames, the data also contains the distances observed in the dated tree.
- MCMC_samples.tar.gz: it is the compressed file containing the tip-to-internode (events and lineage), tip-to-tip MCMC samples in different folders as RData files.
- ev_MCMC_plots.tar.gz: event tip-to-internode distances Bayesian inference autocorrelation and trace plots for both normal and gamma distributions. The plots also contain the probability density function over the histogram of the data, quantile-quantile plots and the cumulative distribution function of the inferred distributions for each event and subsampling.
- lng_MCMC_plots.tar.gz: lineage tip-to-internode distances Bayesian inference autocorrelation and trace plots for both normal and gamma distributions. The plots also contain the probability density function over the histogram of the data, quantile-quantile plots and the cumulative distribution function of the inferred distributions for each lineage node and subsampling.
- t2t_MCMC_plots.tar.gz: tip-to-tip distances Bayesian inference autocorrelation and trace plots for both normal and gamma distributions. The plots also contain the probability density function over the histogram of the data, quantile-quantile plots and the cumulative distribution function of the inferred distributions for each destination tip and subsampling.
